# SPEN is Required for *Xist* Upregulation during Initiation of X Chromosome Inactivation

**DOI:** 10.1101/2020.12.30.424676

**Authors:** Teresa Robert-Finestra, Beatrice F. Tan, Hegias Mira-Bontenbal, Erika Timmers, Cristina Gontan-Pardo, Sarra Merzouk, Benedetto Daniele Giaimo, François Dossin, Wilfred F. J. van IJcken, John W. M. Martens, Tilman Borggrefe, Edith Heard, Joost Gribnau

**Author notes:** These authors contributed equally (Beatrice F. Tan, Hegias Mira-Bontenbal, Erika Timmers).

## Abstract

At initiation of X chromosome inactivation (XCI), *Xist* is monoallelically upregulated from the future inactive X (Xi) chromosome, overcoming repression by its antisense transcript *Tsix*. *Xist* recruits various chromatin remodelers, amongst them SPEN, which are involved in silencing of X-linked genes in *cis* and establishment of the Xi. Here, we show that SPEN plays an important role in the initiation of XCI. *Spen* null female mouse embryonic stem cells (ESCs) are defective in *Xist* upregulation upon differentiation. We find that *Xist*-mediated SPEN recruitment to the Xi chromosome happens very early in XCI, and that SPEN-mediated silencing of the *Tsix* promoter is required for *Xist* upregulation. Accordingly, failed *Xist* upregulation in *Spen*^−/−^ ESCs can be rescued by concomitant removal of *Tsix*. These findings indicate that SPEN is not only required for the establishment of the Xi, but is also crucial in the initiation of the XCI process.

## Introduction

To compensate for gene dosage imbalance between females (XX) and males (XY), female placental mammals randomly inactivate one X chromosome early during embryonic development^1^. In mice, random X chromosome inactivation (XCI) takes place in the epiblast in three phases: initiation, establishment and maintenance. During the initiation phase, the long non-coding RNA (lncRNA) *Xist* is upregulated from the future inactive X (Xi) chromosome^2–4^. *Xist* is located within the X Chromosome Inactivation Centre (*Xic*), an X-linked region required for XCI that contains different *cis*-regulatory elements, including *Tsix*, another lncRNA gene that is transcribed antisense to and completely overlaps *Xist*^5^. *Tsix* negatively regulates *Xist* expression via antisense transcription and chromatin remodelling^6–12^. Together*, Xist* and *Tsix* form a master switch that controls initiation of XCI: in the pluripotent state, *Tsix* is biallelically expressed, repressing both alleles of *Xist*, while upon initiation of XCI the balance changes, resulting in downregulation of *Tsix* and upregulation of *Xist* on the future Xi. *Trans*-regulators including pluripotency factors OCT4, NANOG, REX1 as well as XCI activators are key in controlling this switch by regulating *Xist* and *Tsix* transcription (reviewed in^13^). Later, during the establishment phase of XCI, the 17 kb lncRNA *Xist* spreads in *cis* along the future Xi and recruits different proteins involved in gene silencing that render the X transcriptionally inactive (reviewed in^14^). Active histone marks (H3K4me2/me3, H3 and H4 acetylation) are removed and repressive histone marks are instigated, catalysed by Polycomb group complexes and other protein complexes (reviewed in^15^). Finally, in the maintenance phase, the inactive state of the Xi is epigenetically propagated across cell divisions.

Different studies consisting of *Xist* RNA immunoprecipitations coupled to mass spectrometry^16–18^ and genetic screens^19,20^ identified SPEN (also known as SHARP in human and MINT in mouse) as a crucial factor in the establishment phase of XCI. SPEN is a large protein with four N-terminal RNA Recognition Motifs (RRM) and a highly conserved C-terminal SPOC domain able to recruit different proteins involved in transcriptional silencing^21,22^. SPEN is also involved in the Notch signalling pathway and nuclear receptor signalling, where it acts as a transcriptional corepressor^23,24^.

SPEN is crucial for X-linked gene silencing^16,17,19,20^ by binding the *Xist* Repeat A (RepA) via its RRM domains^17,19^ and interacting via its SPOC domain with the corepressors NCoR/SMRT to recruit/activate histone deacetylase 3 (HDAC3), which is responsible for the removal of histone H3 and H4 acetylation at promoters and enhancers of genes located on the future Xi^16,25^. Despite this crucial role for SPEN in establishment of the Xi, these studies did not report defects in *Xist* upregulation and coating^16,17,19,20,25^ (Supplementary Table 1).

Here, we show that SPEN accumulates on the Xi very early during differentiation and is required for *Xist* upregulation. We show that SPEN has a dual function, required to silence *Tsix* and facilitate *Xist* upregulation, while also stabilizing *Xist* RNA. Together, our results indicate that SPEN is not only necessary for X-linked gene silencing but also plays a crucial earlier role in the regulation of initiation of XCI.

## Results

### SPEN is required for Xist upregulation

Previous work has shown how SPEN is crucial for silencing of X-linked genes, but these studies did not investigate the role of SPEN in initiation of *Xist* expression. Therefore, we generated *Spen* homozygous (*Spen*^−/−^) and heterozygous (*Spen*^+/−^) knockout mouse embryonic stem cells (ESCs) by deleting the complete open reading frame (ORF) using the CRISPR/Cas9 technology (Supplementary Fig. 1a). These lines were generated in a hybrid F1 129/Sv:Cast/EiJ (129/Cast) genetic background with a doxycycline-responsive endogenous *Xist* promoter located on the Cast X chromosome^26^ (Fig. 1a). The S*pen* ORF deletion was verified by PCR on genomic DNA (gDNA) and Western blot analysis (Supplementary Fig. 1b-c). Allele-specific RNA-seq analysis (Supplementary Fig. 1d) of wild type (Wt) undifferentiated ESCs (day 0) containing the doxycycline-responsive *Xist* promoter treated with and without doxycycline for 4 days (Fig. 1b) showed skewed X-linked gene silencing towards the Cast allele (Fig. 1c top-left, Supplementary Fig. 1e). On the other hand, the same analysis in *Spen*^−/−^ ESCs showed impaired X-linked gene silencing (Fig. 1c bottom-left), as described before^16,17,19,20^ (Supplementary Table 1).

**Fig. 1.**
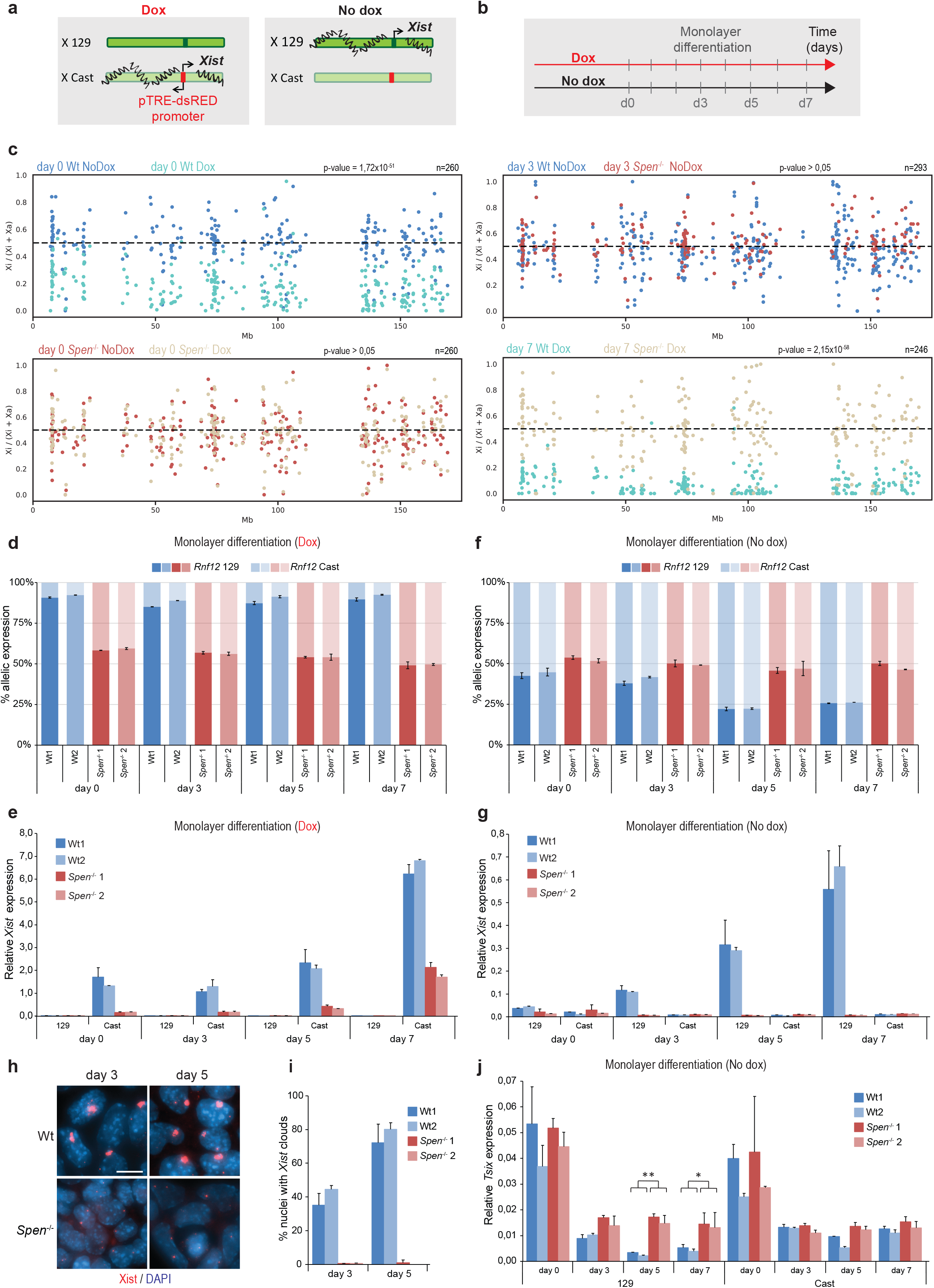
Impaired *Xist* upregulation in *Spen*^−/−^ ESCs upon monolayer differentiation. **a** Overview of the endogenous doxycycline-inducible *Xist* hybrid system used in this study. Addition of doxycycline leads to inactivation of the Cast X chromosome, while no addition of the drug leads to inactivation of the 129 X chromosome. **b** Experimental design to recapitulate XCI. Cells were treated or untreated with doxycycline for 4 days previous to monolayer differentiation. **c** Allelic ratio ((Xi)/(Xi+Xa)) of individual genes along the X chromosome of Wt **(top-left)** and *Spen*^−/−^ **(bottom-left)** undifferentiated (day 0) ESCs with and without doxycycline. Allelic ratio of individual genes along the X chromosome of Wt and *Spen*^−/−^ lines at day 3 of differentiation without doxycycline **(top-right)** and day 7 with doxycycline **(bottom-right)**. Only the genes (n) with sufficient reads in both conditions are shown. Xa = active X chromosome. **d** Percentage of *Rnf12* allelic expression at different time points of monolayer differentiation of two independent Wt and *Spen*^−/−^ ESC lines treated with doxycycline, determined by RT-qPCR. Relative *Rnf12* allelic (129 and Cast) expression was normalized to *Rnf12* total expression and averaged ± SD, n=2 biological replicates. **e** Relative allele-specific *Xist* expression of two independent Wt and *Spen*^−/−^ ESC lines at different time points of monolayer differentiation treated with doxycycline. Average expression ± Standard Deviation (SD), n=2 biological replicates. **f** Same as displayed in **(d)** without doxycycline. **g** Same as displayed in **(e)** without doxycycline. **h** *Xist* RNA FISH (red) of Wt and *Spen*^−/−^ ESC lines at day 3 and 5 of differentiation. Both *Tsix* pinpoints and *Xist* clouds are visible. DNA is stained with DAPI (blue). Scale bar: 10 μm. **i** Quantification of **(h)**, displaying the average percentage of nuclei with *Xist* clouds. Average percentage ± SD, n=2 biological replicates, 150-500 nuclei quantified per replicate. **j** Relative allele-specific *Tsix* expression of Wt and Spen^−/−^ ESCs at different time points of monolayer differentiation, determined by RT-qPCR. Average expression ± SD, n=2 biological replicates. Statistical analysis was done using a two-tailed Student’s t-test comparing four independent replicates per condition (two biological replicates of two different clones). (*) p-value <0.05, (**) p-value <0.01.

To trigger XCI in the context of differentiation, we forced *Xist* upregulation by doxycycline treatment followed by monolayer differentiation (Fig. 1b). Allele-specific RNA-seq analysis of Wt and *Spen*^−/−^ ESCs treated with doxycycline at day 7 of monolayer differentiation (Supplementary Fig. 1d) also revealed a lack of X-linked gene silencing along the entire X-chromosome (Fig. 1c bottom-right, Supplementary Fig. 1e). Similarly, allele-specific RT-qPCR analysis of the X-linked gene *Rnf12* revealed impaired silencing (Fig. 1d). Although previous work indicated that in *Spen*^−/−^ ESCs a group of lowly expressed X-linked genes are susceptible to SPEN-independent gene silencing^27^, our RNA-seq analysis shows no silencing of this specific group of genes (Supplementary Fig. 1f,g). We observed *Xist* upregulation in Wt and *Spen*^−/−^ cells, however doxycycline induction resulted in lower *Xist* expression levels in *Spen*^−/−^ compared to Wt ESCs (Fig. 1e), an effect that was also reported in a recent study^27^.

Next, we recapitulated physiological XCI by monolayer differentiation in the absence of doxycycline to allow normal *Xist* upregulation and X-linked gene silencing (Fig. 1b). Allele-specific RNA-seq analysis of Wt and *Spen*^−/−^ ESCs at day 3 without doxycycline (Supplementary Fig. 1d) shows no significant differences in silencing (Fig. 1c top-right, Supplementary Fig. 1e), making evident the time scale differences between forced *Xist* upregulation in the undifferentiated ESC stage and physiological XCI. Even so, at the latest time points of differentiation (day 5 and 7) *Spen*^−/−^ cells clearly lost the capacity to induce *Rnf12* silencing (Fig. 1f). Remarkably, in *Spen*^−/−^ cells, *Xist* upregulation from the 129 allele was completely abrogated (Fig. 1g), contrasting earlier evidence that suggested that SPEN is not required for *Xist* upregulation and coating 16,17,19,20,25 (Supplementary Table 1). In addition, differentiating *Spen*^−/−^ ESCs lack *Xist* clouds, determined by RNA-FISH (Fig. 1h,i), while *Tsix* was significantly more expressed from the Wt 129 allele in *Spen*^−/−^ cells compared to Wt cells at day 5 and 7 of monolayer differentiation (Fig. 1j), suggesting that SPEN might be necessary for *Tsix* silencing. Importantly, RT-qPCR analysis confirmed proper silencing of pluripotency genes *Rex1* and *Nanog*, and upregulation of endoderm marker *Gata6* (Supplementary Fig. 1h) in *Spen*^−/−^ ESCs upon monolayer differentiation, indicating that loss of *Xist* expression is not related to defective ESCs differentiation.

Furthermore, while *Spen*^+/−^ cells are able to upregulate *Xist* and silence *Rnf12* upon doxycycline treatment followed by monolayer differentiation (Supplementary Fig. 2a,b), they show reduced *Rnf12* silencing upon physiological differentiation without a defect in *Xist* upregulation (Supplementary Fig. 2c,d). Given that SPEN levels are reduced in Spen^+/−^ cells (Supplementary Fig. 1c), these results demonstrate that SPEN dosage is important in XCI.

### Spen rescue leads to normal Xist expression levels

To confirm previous results, we performed a rescue experiment by stably re-expressing *Spen* through introduction of the full-length *Spen* cDNA in the *ROSA26* locus^25^ of *Spen*^−/−^ ESCs (Supplementary Fig. 3a). Successful integration and expression of *Spen* was verified by PCR on gDNA and RT-qPCR, respectively (Supplementary Fig. 3b,c). The generated rescue clones (Clone A, B and C) display two- to three-fold overexpression of *Spen* RNA at the ESC stage and during monolayer differentiation (Supplementary Fig. 3c). In contrast to the *Spen^−/−^* lines, all *Spen*^−/−(cDNA)^ rescue clones express *Xist* at similar levels to Wt clones during monolayer differentiation (Fig. 2a). Allele-specific expression analysis of *Rnf12* and *Tsix* (129 allele) indicated that the silencing defect is partially rescued in *Spen*^−/−(cDNA)^ clones (Supplementary Fig. 3d,e). Interestingly, in undifferentiated *Spen*^−/−(cDNA)^ ESCs clones, *Xist* levels were higher than in Wt controls (Fig. 2a), and *Xist* RNA-FISH analysis revealed a significant percentage of *Spen*^−/−(cDNA)^ ESCs having *Xist* clouds (Fig. 2b,c). This abnormal cloud formation could be related to higher SPEN abundance due to overexpression in the undifferentiated state, possibly stabilizing *Xist* or silencing *Tsix*. In addition, these results indicate that the observed defect in *Xist* expression is SPEN mediated and takes place at the very early initiation steps of XCI.

**Fig. 2.**
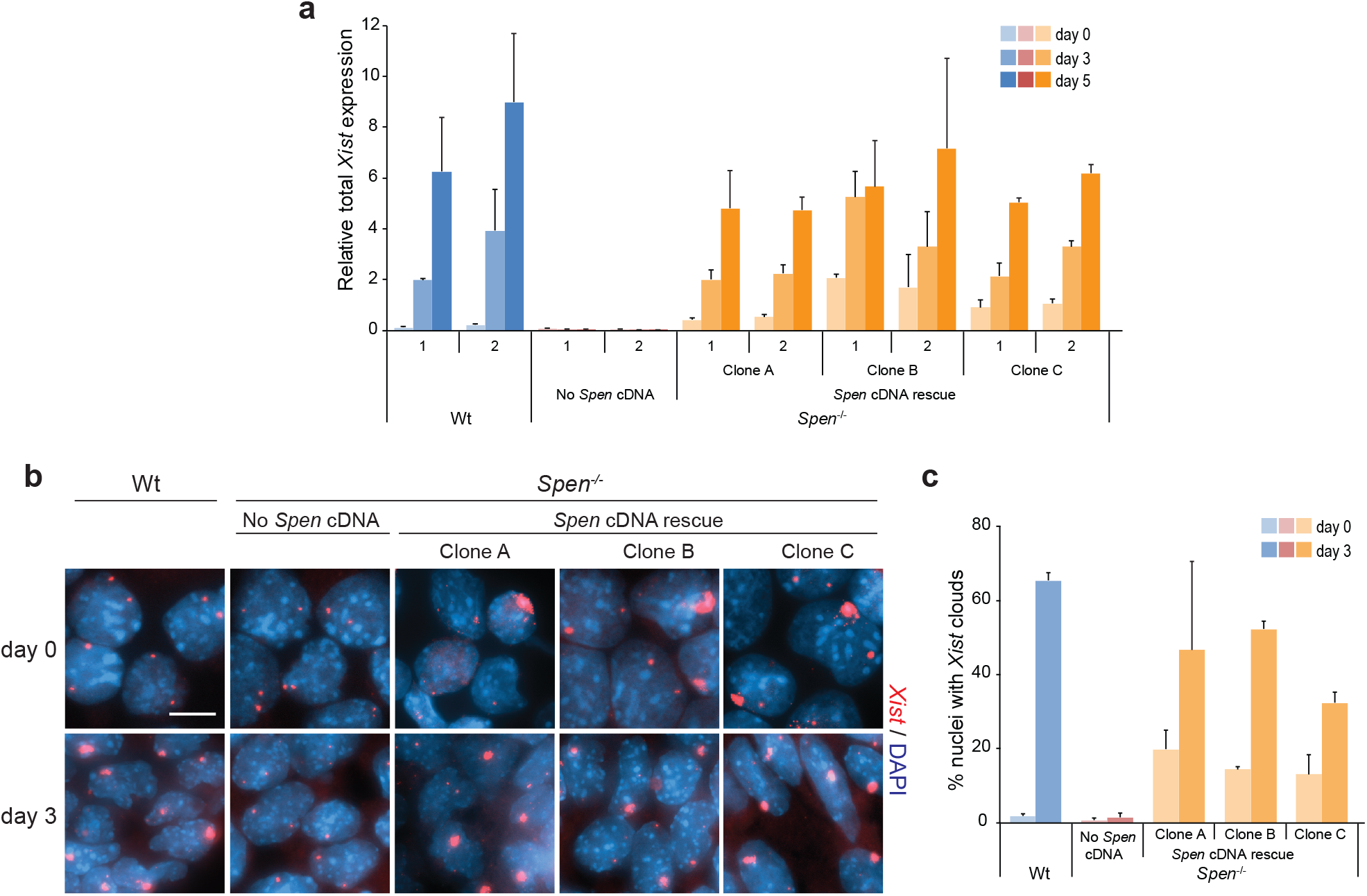
*Spen* cDNA expression in *Spen*^−/−^ ESCs rescues *Xist* expression. **a** Relative total *Xist* expression determined by RT-qPCR at day 0, 3 and 5 of monolayer differentiation of Wt, *Spe*n^−/−^ and three independent rescue clones (Clone A, B and C). Each individual clone was differentiated twice as biological duplicates. Average expression ± SD, n=2 biological replicates. **b** RNA FISH of *Xist* (red) at day 0 and 3 of monolayer differentiation of Wt, *Spen*^−/−^ and *Spen* rescue clones. DNA stained with DAPI (blue). Scale bar: 10 μm. **c** Quantification of **(b)**, showing the percentage of nuclei with *Xist* clouds in each condition. Average percentage ± SD, n=2 biological replicates, 150-400 nuclei quantified per replicate.

**Fig. 3.**
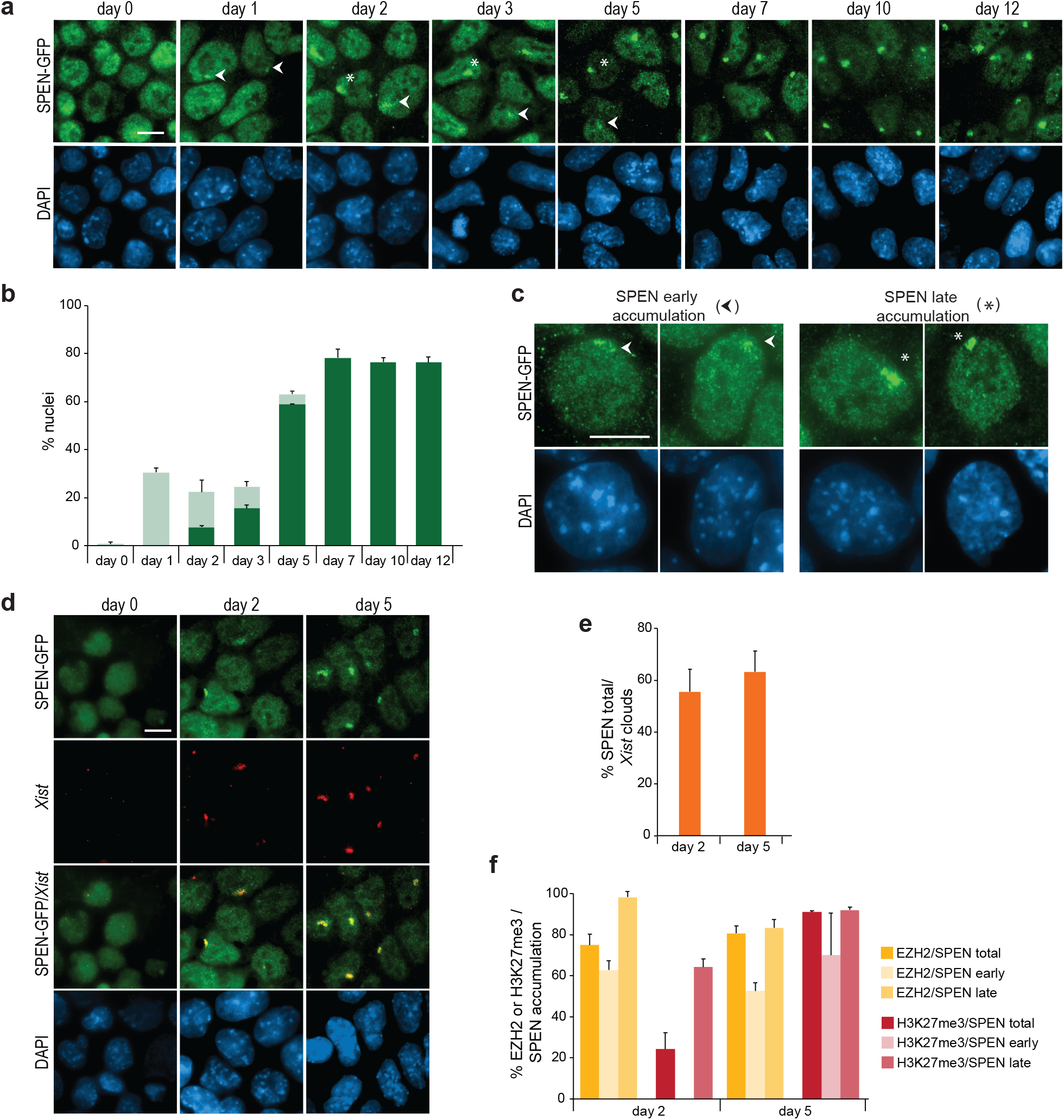
SPEN co-localizes with the Xi chromosome very early during monolayer differentiation. Two different states are detectable: early and late SPEN accumulation on the Xi. **a** SPEN-GFP IF staining (αGFP, green) at different time points of monolayer differentiation (day 0, 1, 2, 3, 5, 7, 10 and 12). White arrowheads mark early SPEN accumulation to the Xi and asterisks late SPEN accumulation. DNA is stained with DAPI (blue). Scale bar: 10 μm. **b** Quantification of **(a)**, depicting the percentage of early and late SPEN accumulation per nuclei at different time points of monolayer differentiation. Average percentage ± SD, n=2 independent clones, 200-500 nuclei quantified per replicate. **c** Examples of early **(left)** and late **(right)** SPEN accumulation (αGFP, green) in differentiating cells. White arrowheads mark early SPEN accumulation to the Xi and asterisks late SPEN accumulation. DNA is stained with DAPI (blue). Scale bar: 10 μm. **d** IF (αGFP, green) combined with *Xist* RNA FISH (red) of SPEN-GFP lines at day 0, 2 and 5 of monolayer differentiation. DNA is stained with DAPI (blue). Scale bar: 10 μm. **e** Percentage of nuclei with a *Xist* cloud that also present SPEN accumulation (total). Average percentage ± SD, n=2 independent clones, 200-500 nuclei quantified per replicate. Calculated from the same data as for **Supplementary Fig. 4d**. **f** Percentage of nuclei with SPEN accumulation (total, early and late) that also present EZH2 accumulation or H3K27me3 pinpoints. Average percentage ± SD, n=2 independent clones, 200-500 nuclei quantified per replicate. Calculated from the same data as for **Supplementary Fig. 4g**.

### SPEN accumulation on the Xi upon ESCs differentiation shows two distinguishable states: early and late SPEN accumulation

In light of the previous results, we expect SPEN to accumulate on the Xi at the very early steps of physiological XCI. To investigate this hypothesis and since commercial SPEN antibodies suitable for immunofluorescence (IF) are lacking, we generated ESCs endogenously expressing a C-terminally tagged SPEN-GFP (Supplementary Fig. 4a). Correct GFP integration was confirmed by PCR on gDNA and by FACS analysis (Supplementary Fig. 4b,c). IF detection of GFP at different time points of monolayer differentiation revealed SPEN accumulation as early as day 1 in about 30% of the nuclei (Fig. 3a,b). Interestingly, we could distinguish two different states: early and late SPEN accumulation on the Xi (Fig. 3a-c). Early accumulation was dispersed and had a lower IF intensity, while SPEN accumulation at later stages was more compact and with a higher IF signal. Early accumulation signals were only detected at the initial days of monolayer differentiation (day 1 to 5), while late accumulation signals appeared at day 2, progressively increased over time and plateaued at day 7 (Fig. 3b).

Previous studies have shown that SPEN and *Xist* co-localize in doxycycline-inducible *Xist* undifferentiated ESC lines^20,25^, although the normal timing and relation between SPEN and *Xist* in differentiating cells remained unexplored. Therefore, we investigated SPEN accumulation in relation to *Xist*, by performing GFP IF combined with *Xist* RNA FISH in differentiating cells (Fig. 3d, Supplementary Fig. 4d). This analysis indicated that at day 2 of differentiation, about 55% of the cells with *Xist* clouds showed SPEN co-localization (Fig. 3e). We also studied the relation of SPEN with other key players in XCI, including the PRC2 catalytic subunit EZH2 and its catalytic product H3K27me3. As found for *Xist*, we could detect co-localization of EZH2 and its associated histone modification with SPEN (Supplementary Fig. 4e-g). At day 2 about 60% of the nuclei with early SPEN accumulation displayed EZH2 accumulation in the absence of H3K27me3 deposition, whereas at day 5 about 90% of the cells showed H3K27me3 enrichment at SPEN positive Xi (Fig. 3f). Our results show that SPEN accumulation on the Xi can be distinguished in an early state and late state of SPEN accumulation, where the early state is marked by small dispersed accumulation of SPEN co-localizing with EZH2 without detectable H3K27me3 at day 2, whereas the late state SPEN accumulation is compact and co-localizes with EZH2 and H3K27me3.

### *Xist* RNA stability is compromised in *Spen*^−/−^ ESCs

We have shown that SPEN is required for endogenous *Xist* upregulation and XCI upon ESCs differentiation. In addition, we noticed that forced *Xist* upregulation resulted in reduced *Xist* RNA levels in *Spen*^−/−^ cells compared to Wt cells (Fig. 1e, Fig. 4a). Accordingly, the percentage of nuclei with *Xist* clouds was lower in *Spen^−/−^* cells with doxycycline induced *Xist* versus Wt cells and *Xist* clouds were in general smaller (Fig. 4b,c). This finding could be explained by a role for SPEN in stabilizing *Xist* RNA by complex formation. To study the role of SPEN in *Xist* stability, we determined the half-life of doxycycline-induced *Xist* RNA in *Spen*^−/−^ and Wt ESCs treated with actinomycin D to block its transcription. The remaining levels of *Xist* RNA at different time-points was assessed by RT-qPCR, and the *Xist* RNA decay rate and half-life were then calculated (Fig. 4d). In Wt ESCs, the *Xist* RNA half-life was 6h 37min, similar to what was described in a previous study^28^, whereas the *Xist* half-life was reduced to 3h 52min in *Spen*^−/−^ cells. These results indicate that SPEN plays a role in promoting *Xist* RNA stability. However, these results cannot explain why physiological *Xist* upregulation is lost in *Spen^−/−^* ESCs upon differentiation.

**Fig. 4.**
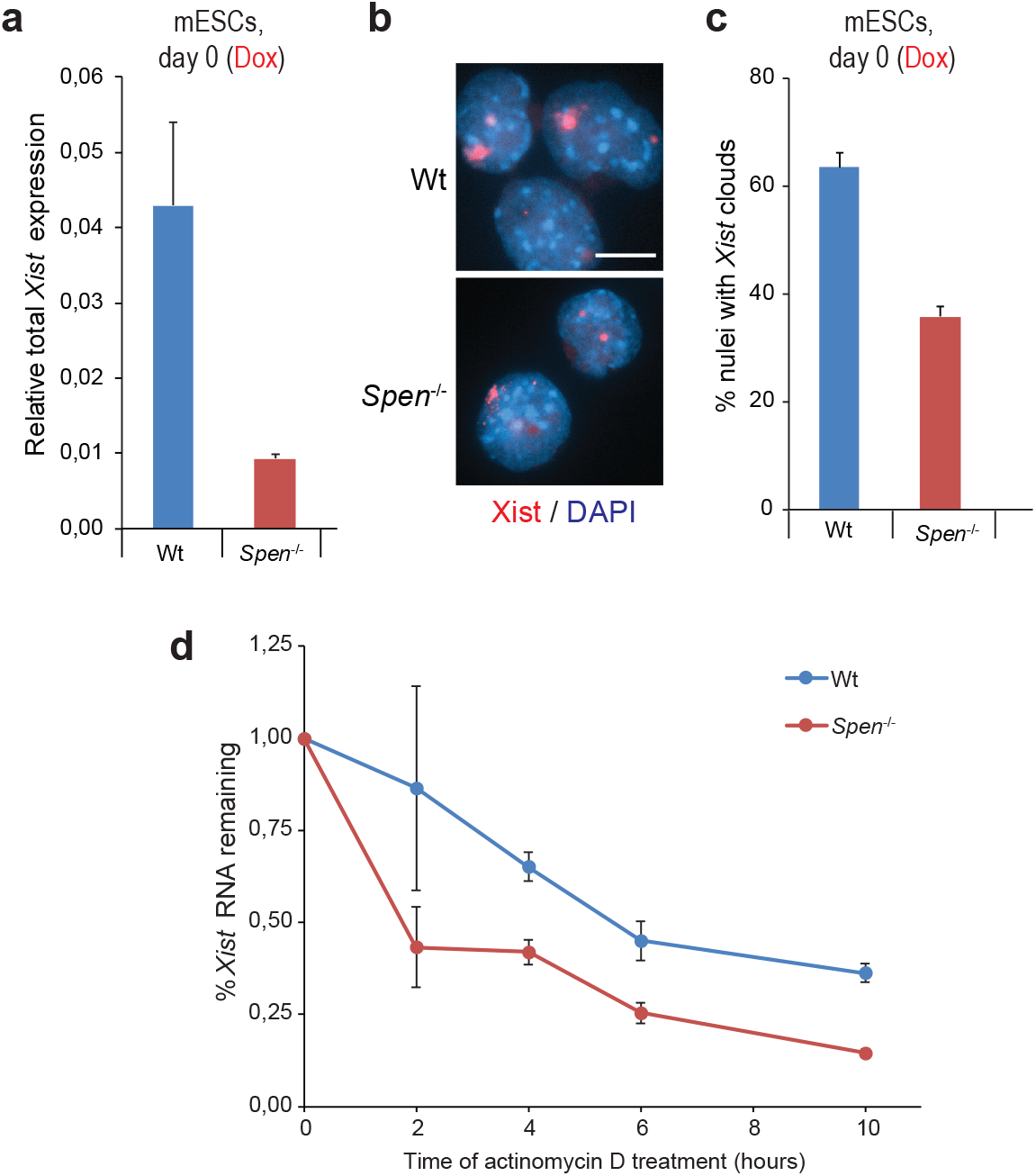
SPEN plays a role in *Xist* RNA stability. **a** Relative total *Xist* expression in undifferentiated Wt and *Spen*^−/−^ ESCs upon *Xist* induction with doxycycline for 4 days. Average expression ± SD, n=2 biological replicates. **b** *Xist* RNA FISH (red) of undifferentiated Wt and *Spen*^−/−^ ESCs treated with doxycycline to induce *Xist* expression. DNA is stained with DAPI (blue). Scale bar: 10 μm. **c** Quantification of **(b)**, displaying the percentage of nuclei with *Xist* clouds. Average percentage ± SD, n=2 biological replicates, 100-150 nuclei quantified per replicate. **d** Percentage of *Xist* RNA remaining at different time points of actinomycin D treatment (t = 0, 2, 4, 6 and 10 hours) in undifferentiated Wt and *Spen*^−/−^ ESCs treated with doxycycline. Normalized relative expression to t=0 ± SD, n=2 biological replicates.

### SPEN, HDAC3 and H3K27ac are enriched at the Tsix regulatory region

Our results showed that at late days of monolayer differentiation, *Spen*^−/−^ cells display higher *Tsix* levels than control cells (Fig. 1j), suggesting that SPEN might be recruited by *Xist* to silence *Tsix*. Hence, we explored SPEN genomic binding at the *Xist-Tsix* locus using published SPEN CUT&RUN data in undifferentiated ESCs with a doxycycline-responsive *Xist* promoter^25^. SPEN accumulates on the *Xist* gene body as well as on the *Tsix* regulatory region, SPEN accumulation is evident at 24 hours of doxycycline induction, but more prominent after 4 and 8 hours of induction (Fig. 5a,b). This *Tsix* regulatory region comprises the minor and major *Tsix* promoters and *Xite*, an enhancer of *Tsix*^29^. *Xist-*mediated recruitment of SPEN is important for recruitment and/or activation of HDAC3, responsible for the removal of H3K27ac from the future Xi^16,30^. Analysis of published HDAC3 and H3K27ac ChIP-seq data from undifferentiated female ESCs upon 24h *Xist* induction^30^ reveals HDAC3 binding and H3K27ac loss at the *Tsix* regulatory region, where SPEN is recruited (Fig. 5a). Moreover, X-linked *Rnf12* (Supplementary Fig. 5a) and *Pgk1* (Supplementary Fig. 5b) display SPEN and HDAC3 enrichment at their promoters and H3K27ac loss upon *Xist* induction. Interestingly, *Rnf12*, an early silenced gene^31^, shows SPEN and HDAC3 promoter binding in the undifferentiated state without *Xist* induction, suggesting that in the undifferentiated state *Xist* might be sufficiently expressed at very low levels to partly silence *Rnf12*, explaining the *Rnf12* allelic ratio difference between Wt and *Spen*^−/−^ ESCs at day 0 (Fig. 1f, Supplementary Fig. 3d).

**Fig. 5.**
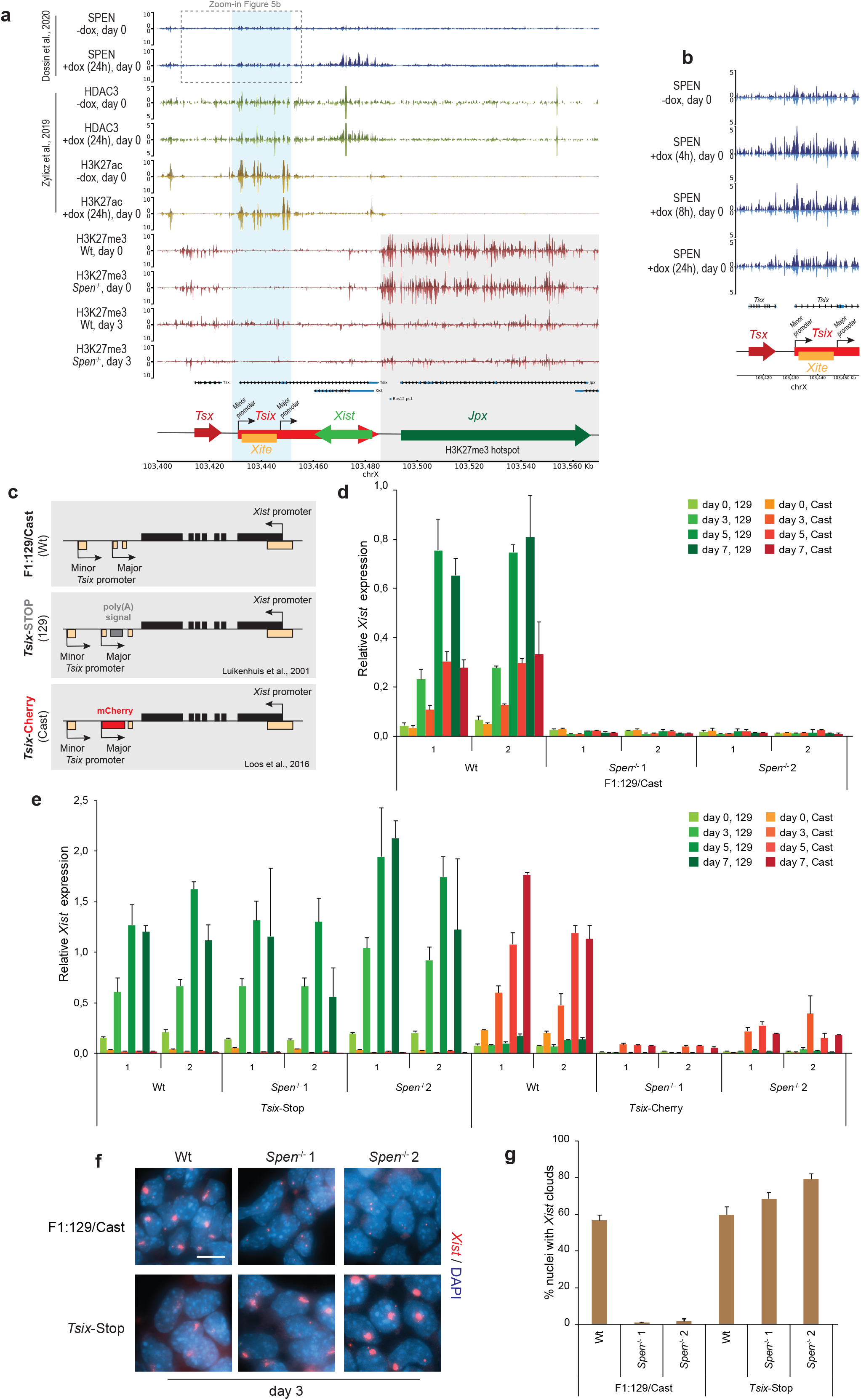
*Tsix* regulatory region shows SPEN, HDAC3 and H3K27ac enrichment. SPEN is required to silence *Tsix*, to allow *Xist* upregulation upon XCI initiation. **a** Genome browser tracks showing allele-specific SPEN, HDAC3, H3K27ac and H3K27me3 binding at the *Xist*-*Tsix* locus. The top part (dark) of each track represents the Xi and the bottom (light) the Xa. SPEN CUT&RUN **(blue, top)** profile in *Xist* inducible undifferentiated ESCs (day 0) untreated or treated with doxycycline (24h)^25^. HDAC3 **(green, middle-top)** and H3K27ac ChIP-seq **(yellow, middle-bottom)** in *Xist* inducible undifferentiated ESC (day 0) untreated or treated with doxycycline (24h)^30^. H3K27me3 ChIP-seq **(red, bottom)** in Wt and *Spen*^−/−^ ESCs at day 0 and 3 of monolayer differentiation. The light-blue rectangle highlights the *Tsix* regulatory region, including the *Tsix* minor and major promoters, and *Xite*. The light-gray rectangle highlights the H3K27me3 hotspot region. The dashed rectangle indicates the genomic region in **(b)**. **b** Zoom-in view of the *Tsix* promoter region, showing the SPEN CUT&RUN profile in undifferentiated ESCs (day 0) untreated or treated (4h, 8h and 24h) with doxycycline. **c** Schematic overview of the *Tsix-*defective ESC lines used to study the role of SPEN in *Tsix* silencing and *Xist* upregulation, namely, Wt (F1:129/Cast) **(top)**, *Tsix-*Stop **(middle)** and *Tsix-*Cherry **(bottom)**. The *Tsix-* Stop line is defective for *Tsix* in the 129 allele; the *Tsix-*Cherry line is defective in the Cast allele. Homozygous deletion of *Spen* was performed in the three lines. **d** Relative allele-specific *Xist* expression in Wt and *Spen*^−/−^ F1:129/Cast ESC lines at different time points of monolayer differentiation. Each individual clone was differentiated twice in biological duplicates. Average expression ± SD, n=2 biological replicates. **e** Same as in **(d)** for Wt and *Spen*^−/−^ *Tsix-*Stop and *Tsix-*Cherry ESC lines. **f** *Xist* RNA FISH (red) at day 3 of monolayer differentiation in F1:129/Cast and *Tsix-*Stop Wt and *Spen*^−/−^ ESC lines. DNA is stained with DAPI (blue). Scale bar: 10 μm. **g** Quantification of **(f)**, showing the percentage of nuclei with *Xist* clouds. Average percentage ± SD, n=2 biological replicates, 150-500 nuclei quantified per replicate.

To test whether SPEN recruitment leads to H3K27me3 accumulation, we performed H3K27me3 ChIP-seq on *Spen*^−/−^ and Wt ESCs prior to and after differentiation (day 3) (Fig. 5a). This analysis revealed H3K27me3 enrichment in the *Tsix* regulatory region in Wt cells upon differentiation, while *Spen*^−/−^ cells do not show H3K27me3 enrichment in accordance with the lack of *Xist* upregulation. Interestingly, the H3K27me3 hotspot, located at the 3’ end of *Tsix*^32,33^ is clearly reduced upon differentiation both in Wt and *Spen*^−/−^ cells, while Wt and *Spen*^−/−^ cells show no difference in H3K27me3 levels, indicating that SPEN does not play a role in H3K27me3 enrichment at the hotspot (Fig. 5a). Altogether, these data support a model where *Xist-*mediated SPEN recruitment leads to *Tsix* promoter silencing.

### SPEN is required to silence Tsix to allow Xist upregulation

To further investigate the role of SPEN in silencing *Tsix*, we generated compound *Spen*^−/−^:*Tsix* defective hybrid ESC lines. If *Xist*-mediated recruitment of SPEN to the *Tsix* regulatory region is crucial for *Tsix* silencing and *Xist* upregulation, we expect *Xist* upregulation upon differentiation to be rescued in these double knockout cell lines. We made use of a *Tsix-*Stop line containing a triple poly(A) signal downstream of the major *Tsix* promoter that blocks its transcription on the 129 allele^7^, and a *Tsix-*Cherry line with a mCherry coding sequence introduced downstream of the major *Tsix* promoter on the Cast allele^34^ (Fig. 5c). As controls, we generated *Spen*^−/−^ ESCs in the same hybrid background (F1:129/Cast) where both *Xist* alleles are intact^35^, in contrast to the previously studied ESCs that contained one doxycycline-inducible *Xist* allele. Deletion of the *Spen* ORF was confirmed in all cell lines by PCR on gDNA and Western blot analysis (Supplementary Fig. 6a,b). For each line we generated two independent knockout clones. All the ESC lines were differentiated in parallel and in biological duplicates, followed by allele-specific *Xist* RNA expression analysis by RT-qPCR. As expected, the F1:129/Cast control *Spen*^−/−^ line showed no *Xist* upregulation (Fig. 5d), similar to the phenotype observed in the *Spen* knockout clones generated in the heterozygous doxycycline-inducible *Xist* cell line (Fig. 1g). Remarkably, the introduction of a poly(A) signal in *Spen*^−/−^:*Tsix-*Stop lines fully rescued *Xist* upregulation upon differentiation (Fig. 5e), indicating that SPEN is required for *Xist* upregulation via *Tsix* repression. However, the *Spen^−/−^:Tsix-*Cherry lines displayed only mild upregulation of *Xist* upon differentiation (Fig. 5e). These differences in the rescue phenotype between *Spen^−/−^:Tsix-*Cherry and *Spen*^−/−^:*Tsix-*Stop lines can be explained by remaining *Tsix* transcription in the *Tsix-*Cherry line, which is fully ablated in the *Tsix-*Stop lines (Supplementary Fig. 6c). As expected, the *Spen*^−/−^: *Tsix-*Stop lines are not able to silence X-linked *Rnf12* despite normal *Xist* levels in the *Tsix-*Stop line (Supplementary Fig. 6d). Moreover, the Spen^−/−^:*Tsix-*Stop lines display *Xist* clouds that were similar in morphology (Fig. 5f) and number compared to Wt F1:129/Cast cells (Fig. 5g).

Taken together, our results indicate a novel and essential role for SPEN in *Xist* upregulation, mainly via silencing of *Tsix*, and to a lesser extent by stabilizing *Xist* RNA. SPEN is therefore not only crucial in X-linked gene silencing but also in the early initiation steps of XCI, playing a role in the feedforward loop leading to *Xist* activation (Fig. 6).

**Fig. 6.**
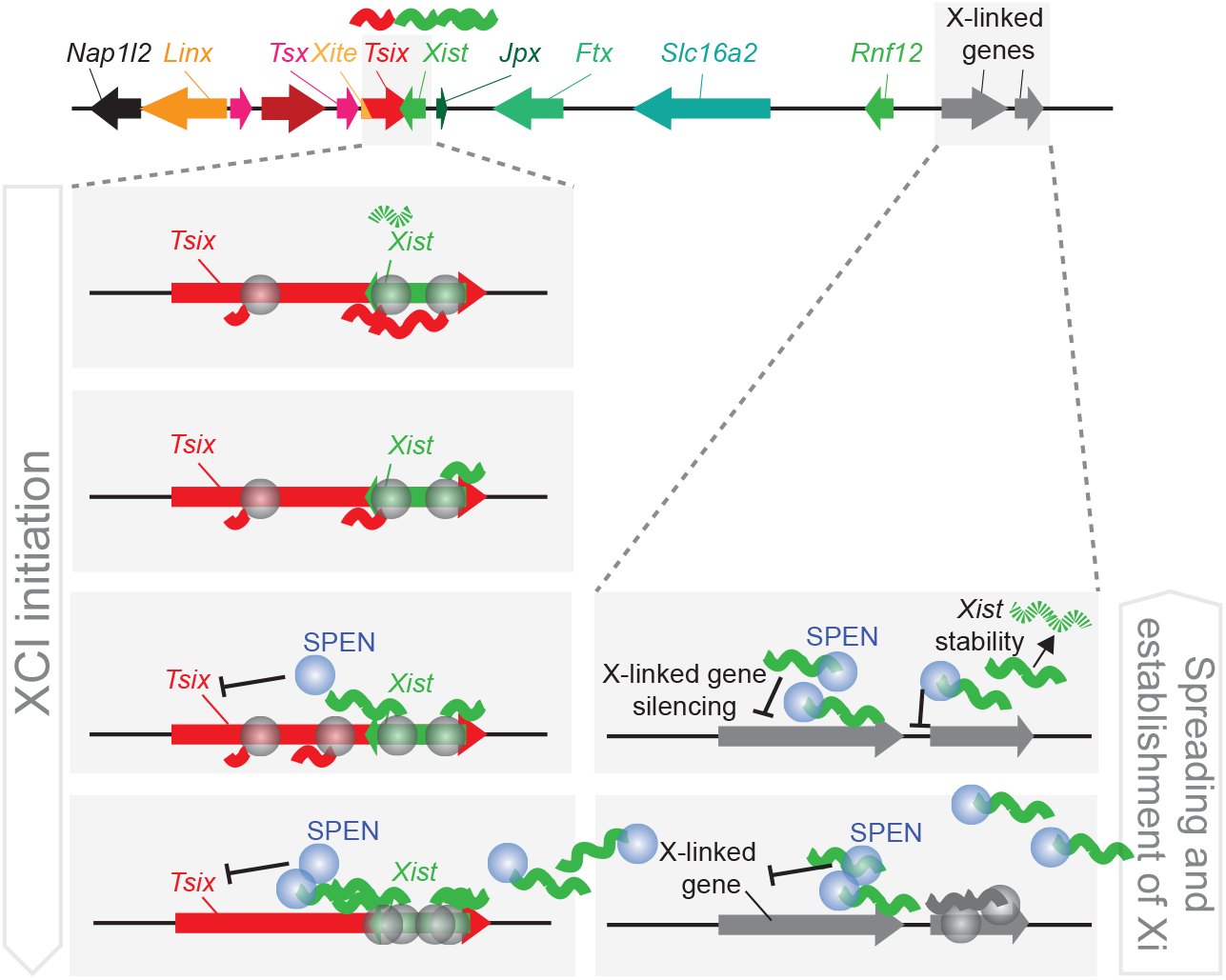
Model of the role of SPEN in XCI. In the undifferentiated state, *Tsix* transcription represses *Xist* and *Tsix* RNA levels are high in ESCs. Upon differentiation, at the start of XCI, *Xist* transcription increases, the nascent *Xist* transcripts recruit SPEN that is necessary to silence the *Tsix* promoter via removal of active H3K27ac marks from its promoter. Consequently, the silencing of *Tsix* allows *Xist* upregulation, accumulation and spreading. The spreading of SPEN-*Xist* along the X chromosome allows the silencing of X-linked genes. SPEN also plays a role in *Xist* stability. This model, in addition to previous evidence, proposes that SPEN is not only key in X-linked gene silencing, but also in initiation of XCI. In the figure, the blue circles depict SPEN and the grey circles RNA Polymerase II.

## Discussion

The role of the silencing factor SPEN in the establishment phase of XCI has been addressed in various studies, but whether SPEN is relevant for the initiation phase, involving *Xist* upregulation, remains unknown. Here, we show that SPEN-defective ESCs do not upregulate *Xist* upon differentiation and study the molecular mechanism behind this observation. To explore the role of SPEN in XCI, various studies used *Xist-*inducible ESC lines to generate SPEN knockdown or knockout cell lines (Supplementary Table 1). Forcing *Xist* expression in ESCs is a powerful way to understand X-linked gene silencing, but is not suitable to investigate the initiation phase of XCI. Studies exploring the role of SPEN in cells undergoing physiological XCI involved *Spen* knockdown strategies^17,20^, therefore, the levels of SPEN might have been sufficient to allow normal *Xist* upregulation, while showing a defect in X-linked gene silencing. Likewise, our *Spen*^+/−^ ESC lines show *Xist* upregulation, but reduced X-linked silencing compared to Wt lines. In agreement with our results, one study reports lower *Xist* abundance and cloud formation upon forced *Xist* induction from the endogenous locus in *Spen*^−/−^ ESCs^27^. Another recent study suggested that *Spen*^−/−^ ESCs are not able to differentiate upon Leukemia Inhibitory Factor (LIF) removal and differentiation towards neural progenitor cells^36^, while we observe that *Spen*^−/−^ ESCs display a normal morphology in the undifferentiated state and undergo normal differentiation upon monolayer differentiation. Nevertheless, we observe more cell death of *Spen*^−/−^ cells compared to Wt cells upon monolayer differentiation, although we consider this might not be XCI-related since X0 *Spen*^−/−^ cells also die upon differentiation (data not shown). This observation is probably not surprising since SPEN plays a role in various biological processes as a transcriptional repressor^23,24,37^.

Previous studies performed with doxycycline-inducible *Xist* lines show that SPEN and *Xist* co-localize soon after *Xist* induction^20,25^. However, *Xist* upregulation from a doxycycline-inducible promoter happens at a very different timescale than during physiological XCI. In the present study, we observe that upon *Spen*-GFP ESCs monolayer differentiation we can distinguish early and late SPEN accumulation on the Xi. Cells with SPEN early accumulation are detectable from day 1 of differentiation with their number decreasing overtime, while the number of cells with SPEN late accumulations progressively increases. Other XCI key players, such EZH2 and H3K27me3, also sequentially accumulate on the Xi.

In addition, we provide evidence that SPEN is required to silence *Tsix* to allow *Xist* upregulation. (I) In *Spen*^−/−^ cells, we detect higher levels of *Tsix* at the latest time points of monolayer differentiation, compared to Wt cells. In line with this, paternal *Tsix* levels in female E3.5 blastocysts with a mutated *Xist* RepA, necessary for SPEN recruitment, are higher than in Wt blastocysts^38^, suggesting that higher *Tsix* levels in *Spen*^−/−^ cells are due to defective *Tsix* silencing, rather than a lack of *Xist* antisense transcription. (II) SPEN overexpression in ESCs leads to higher *Xist* levels and ectopic *Xist* cloud formation in the undifferentiated state, suggesting that SPEN overexpression might lead to partial silencing of the *Tsix* promoter, facilitating *Xist* expression. (III) SPEN binds to active promoters and enhancers to silence the X chromosome^25^. Spatial proximity to the *Xist* locus is a strong predictor of X-linked gene silencing efficiency^26,39^. One of the closest actively transcribed promoters to *Xist* is *Tsix* and we indeed observe SPEN binding at the *Tsix* regulatory region. (IV) *Spen*^−/−^ cell lines cannot upregulate *Xist*, while compound *Spen*:*Tsix* defective cell lines (*Spen*^−/−^:*Tsix*-Stop) display normal *Xist* levels upon differentiation, indicating that in the absence of SPEN, *Tsix,* which acts a as brake on *Xist* transcription, cannot be silenced. Furthermore, our results show that SPEN plays a role in *Xist* stability, however, we also observe that *Xist* levels in differentiating *Spen*^−/−^:*Tsix*-Stop cells are comparable to Wt cells, suggesting that *Xist* stability might not be compromised. This difference may be attributed to the massive *Xist* overexpression in *Xist-*inducible systems, where the excess of *Xist* RNA molecules, in the absence of SPEN, may affect overall *Xist* RNA stability readings.

During initiation of XCI, SPEN helps remodel the chromatin environment of the *Xist-Tsix* locus. SPEN and HDAC3 are present at the *Tsix* regulatory region and H3K27ac levels decrease upon *Xist* induction. Accordingly, we propose that during XCI initiation, SPEN binds *Xist* nascent transcripts^25^, recruits and/or activates HDAC3^16,30^, which removes histone acetylation marks, weakening *Tsix* promoter activity and facilitating *Xist* expression (Fig. 6).

The mechanism driving the symmetry breaking event leading to monoallelic upregulation of *Xist* has been a focus of many studies. Several studies showed that the *Xic* and more specifically the *Xist*-*Tsix* master switch are tightly regulated in a deterministic process, involving monoallelic down-regulation of *Tsix* or X-pairing mechanisms^40–42^. Our studies, and studies of others, indicate that a gene regulatory network composed of trans-acting activators and inhibitors of XCI, involving positive and negative feedback loops, instruct the *cis*-regulatory landscape of the *Xic* to direct monoallelic upregulation of *Xist*^12,43–45^. Our present work reveals that *Tsix* silencing not only involves downregulation of inhibitors of XCI, including pluripotency factors OCT4, SOX2, KLF4, c-MYC and REX1^46–48^, but also involves *Xist-*mediated recruitment of SPEN. This can happen on either allele, but asynchronous *Xist* transcription bursts will facilitate *Xist*-mediated monoallelic silencing of *Tsix* through SPEN, resulting in further upregulation of *Xist* and concomitant silencing of the XCI activator *Rnf12*, providing a negative feedback loop to prevent upregulation of *Xist* on the future active X chromosome.

## Acknowledgements

We thank Anniek Meesters and Esther Sleddens-Linkels for technical help; Kristian Helin for providing the EZH2 antibody; Martine M. Jaegle for sharing with us a floxed-puromycin resistance cassette and all the members of the Erasmus MC Developmental Biology department for useful discussions.

T.R.F. is supported by an Erasmus MC grant (Mrace). B.F.T., H.M.B., E.T. and J.G. are supported by the Oncode Institute. T.B. is supported by the Deutsche Forschungsgemeinschaft (DFG, German Research Foundation) - TRR81-A12 and BO 1639/9-1, the Behring-Röntgen foundation and Excellence Cluster for Cardio Pulmonary System (ECCPS) in Giessen. B.D.G. is supported by a research grant of the University Medical Center Giessen and Marburg (UKGM) and by a Prize of the Justus Liebig University Giessen”.

## Author contributions

T.R.F. and J.G. conceived the project and designed the experiments. T.R.F. and E.T. performed most of the experimental work and data analysis. H.M.B. performed the ChIP-seq experiments. B.F.T. performed all the bioinformatic analysis. C.G.P. helped setup the high molecular weight western blot analysis. S.M. setup the allele-specific RT-qPCR analysis. B.D.G., F.D., T.B. and E.H. provided valuable resources. H.M.B. and J.G. supervised the work. T.R.F. and J.G. wrote the manuscript with input from all the authors. H.M.B. helped review and edit the manuscript. J.W.M.M. and J.G. were responsible for the founding acquisition. RNA-seq and ChIP-seq datasets were generated in the Erasmus MC Center for Biomics led by W.v.IJ.

## Supplementary figure legends

**Supplementary Fig 1.**
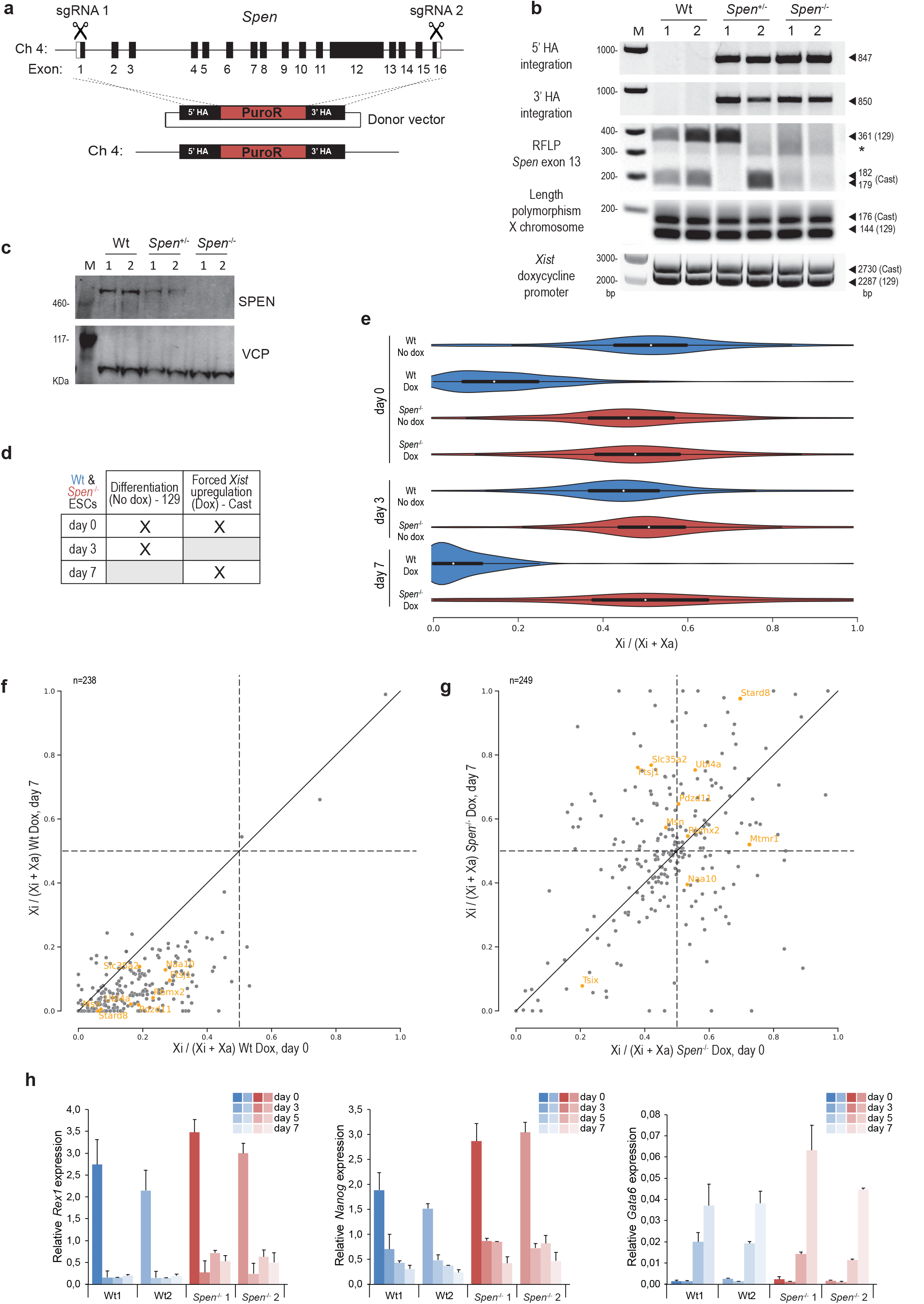
Generation and characterization of *Spen*^+/−^ and *Spen*^−/−^ ESC lines. RNA-seq analysis of Wt and *Spen*^−/−^ ESCs. | Related to **Fig. 1**. **a** Targeting strategy used to generate *Spen* knockout ESC lines using the CRISPR/Cas9 system. Two single guide RNAs (sgRNA) targeting the 5’ and 3’ region of the *Spen* ORF were used to integrate a Puromycin resistance (PuroR) cassette. **b** PCR genotyping to identify optimal heterozygote (^+/−^) and homozygote (^−/−^) *Spen* knockout ESC clones. Specific 5’ and 3’ integration of the PuroR cassette in the *Spen* locus. Restriction Fragment Length Polymorphism (RFLP) analysis on *Spen* exon 13 to determine the absence of *Spen* ORF in one or both alleles. Verification of the presence of two X chromosomes per ESC line. Genotyping of the *Xist* endogenous doxycycline-inducible promoter. M = DNA ladder. (*) = unspecific band. **c** SPEN western blot of two independent Wt, *Spen*^+/−^ and *Spen*^−/−^ ESC clones. VCP was used as a loading control. M = High molecular weight protein ladder. **d** Overview of the RNA-seq libraries generated in this study, summarizing the experimental conditions performed in Wt and *Spen*^−/−^ ESCs at different time points of monolayer differentiation (day 0, 3 and 7), treated with or without doxycycline (X in the panel). Each condition includes two biological replicates, adding up to a total of 16 RNA-seq libraries. **e** Violin plots depicting the distribution of the allelic ratios ((Xi)/(Xi+Xa)) of X-linked genes in Wt and Spen^−/−^ ESCs, untreated or treated with doxycycline at days 0, 3 and 7 of differentiation. This figure summarizes data in **Fig. 1c**. The box plots inside the violin plots show the median and interquartile range. **f** Scatter plot showing the allelic ratio (Xi/(Xi+Xa)) of X-linked genes in Wt ESCs treated with doxycycline at day 0 (x-axis) and day 7 (y-axis) of differentiation. Highlighted in orange are the lowly silenced genes in *Spen*^−/−^ ESC previously identified^27^. Only the genes (n) with sufficient reads in both conditions are shown. Dashed lines = allelic ratio of 0.5; Diagonal solid line = equal ratio in both conditions. **g** Same as in **(f)**, for *Spen*^−/−^ ESCs. **h** *Rex1*, *Nanog* and *Gata6* relative expression of Wt and *Spen*^−/−^ ESC lines upon monolayer differentiation not treated with doxycycline. Average expression ± SD, n=2 biological replicates.

**Supplementary Fig 2.**
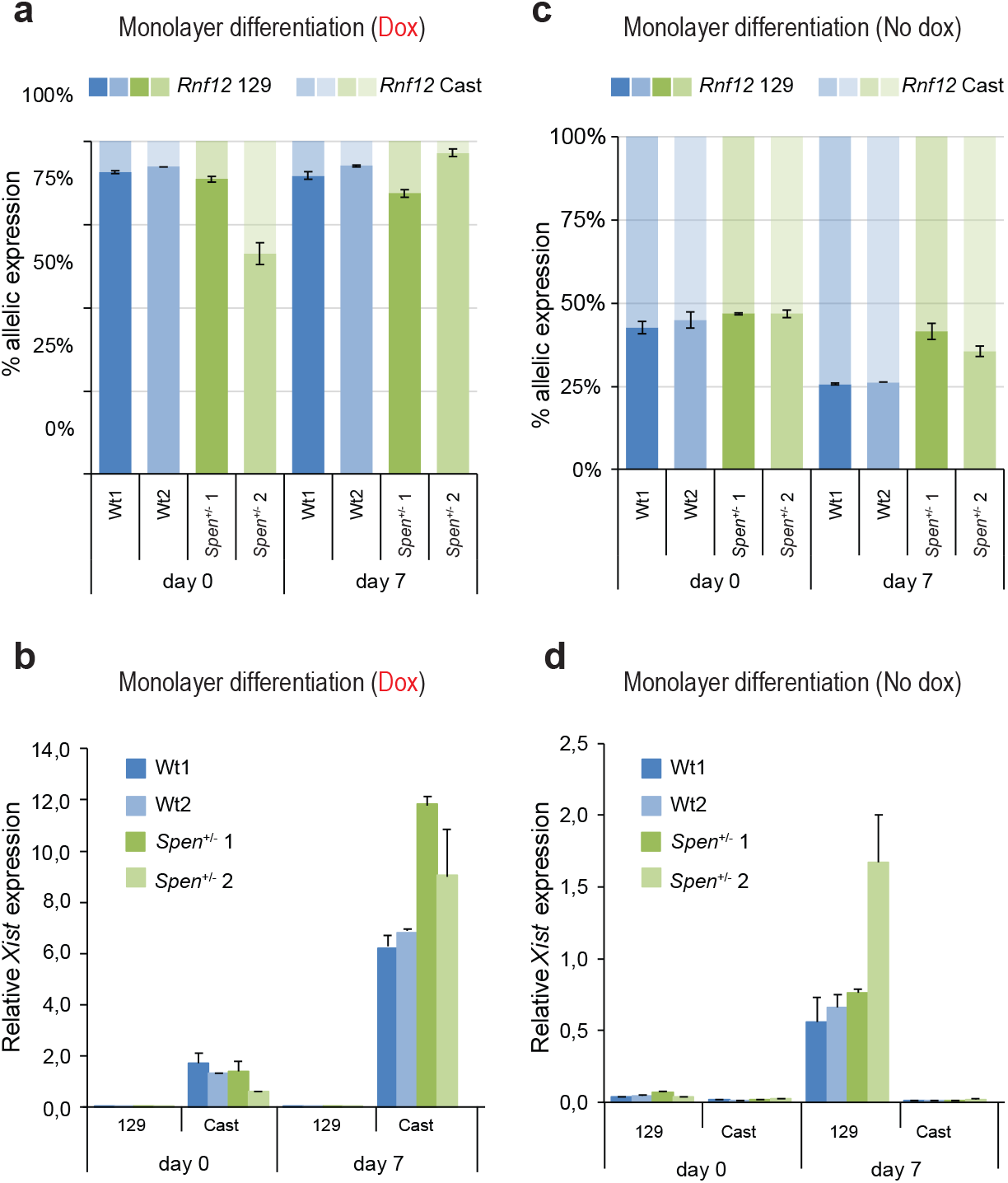
*Spen*^+/−^ ESCs lines are able to upregulate *Xist*, but show less X-linked gene silencing, compared to Wt ESCs upon monolayer differentiation. | Related to **Fig. 1**. **a** *Rnf12* percentage of allelic expression of Wt and *Spen*^+/−^ lines treated with doxycycline at day 0 and 7 of differentiation. Wt samples are the same as in **Fig. 1d**. Relative *Rnf12* allelic (129 and Cast) expression was normalized to *Rnf12* total expression and averaged ± SD, n=2 biological replicates. **b** Relative allele-specific *Xist* expression of two Wt and *Spen*^+/−^ ESC clones treated with doxycycline at day 0 and 7 of monolayer differentiation, determined by RT-qPCR. Wt samples are the same as in **Fig. 1e**. Average expression ± SD, n=2 biological replicates. **c** Same as displayed in **(a)** without doxycycline. Wt samples are the same as in **Fig. 1f**. **d** Same as displayed in **(b)** without doxycycline. Wt samples are the same as in **Fig. 1g**.

**Supplementary Fig 3.**
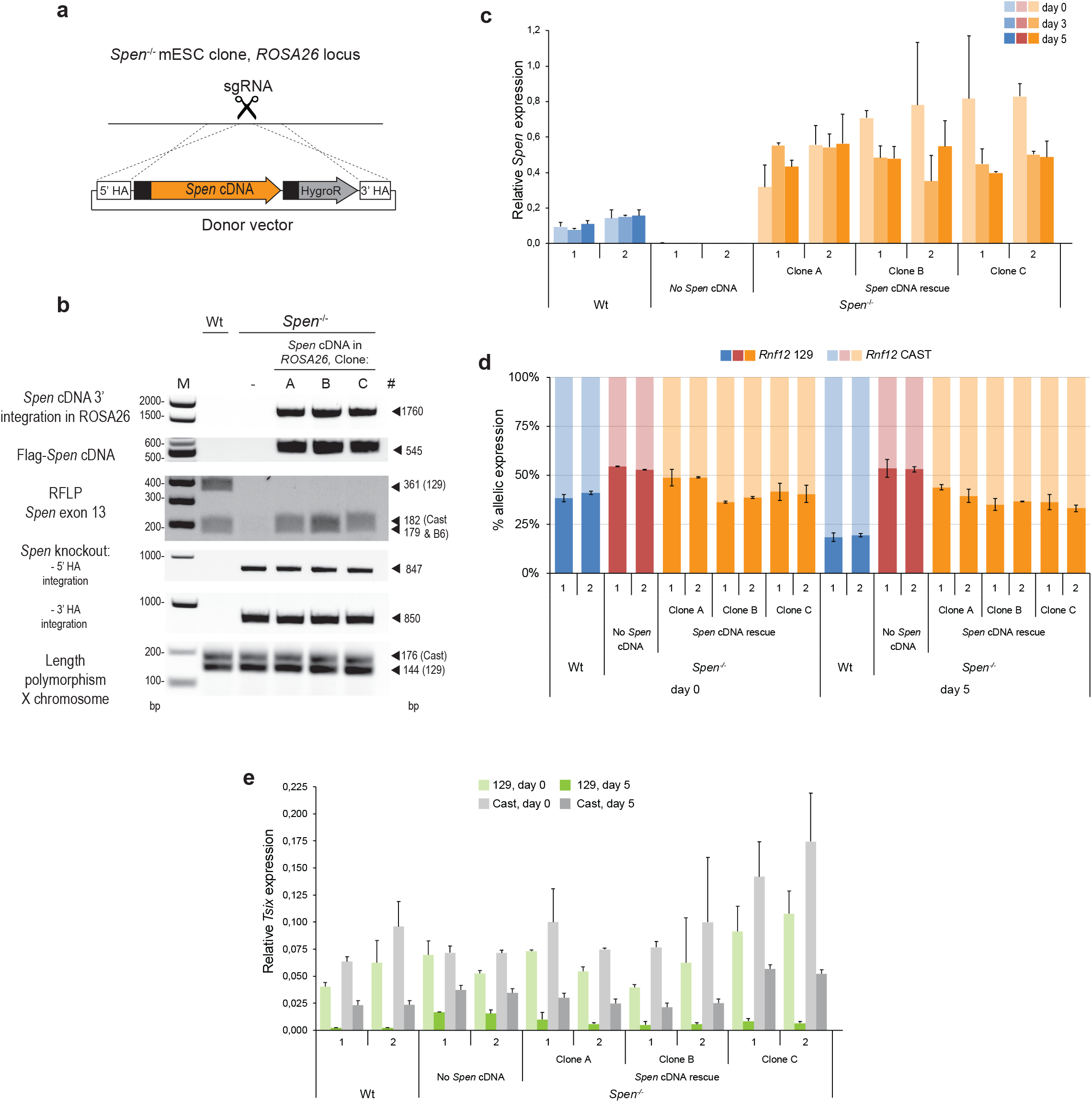
*Spen* cDNA rescue ESC lines characterization. |Related to **Fig. 2**. **a** Schematic overview of the generation of *Spen*^−/−^ ESC lines able to stably express the *Spen* cDNA from the *ROSA26* locus. These lines were made using a sgRNA targeting the *ROSA26* locus and a vector coding for the *Spen* cDNA and a hygromycin resistance cassette (HygroR)^25^. **b** Strategy to identify correct *Spen* cDNA rescue clones by PCR on gDNA. Specific 3’ *Spen* cDNA integration in the *ROSA26* locus. Primer on the Flag-tag present in the *5’ Spen* cDNA end to identify those clones containing the *Spen* cDNA vector. Specific 5’ and 3’ integration of the PuroR cassette in the *Spen* locus to identify the *Spen*^−/−^ line. RFLP analysis on *Spen* exon 13 to determine the presence of *Spen* in the *ROSA26* locus and its absence in the *Spen*^−/−^ cells; the *Spen* rescue cDNA sequence is C57BL/6 (B6). Verification of the presence of two X chromosomes per ESC line, making use of a length polymorphism on the X chromosome. M = DNA ladder. **c** Relative *Spen* expression in Wt, *Spen*^−/−^ and *Spen* cDNA rescue ESC lines (Clone A, B and C) at day 0, 3 and 5 of monolayer differentiation. Each individual clone was differentiated twice in biological duplicates. Average expression ± SD, n=2 biological replicates. **d** Percentage of *Rnf12* allelic expression at day 0, 3 and 5 of monolayer differentiation of Wt, *Spe*n^−/−^ and three *Spen* cDNA rescue clones (Clone A, B and C), determined by RT-qPCR. Relative *Rnf12* allelic (129 and Cast) expression was normalized to *Rnf12* total expression and averaged ± SD, n=2 biological replicates. **e** Relative allele-specific *Tsix* expression at day 0 and 5 of monolayer differentiation of Wt, *Spe*n^−/−^ and three *Spen* cDNA rescue clones (Clone A, B and C), determined by RT-qPCR. Average expression ± SD, n=2 biological replicates.

**Supplementary Fig 4.**
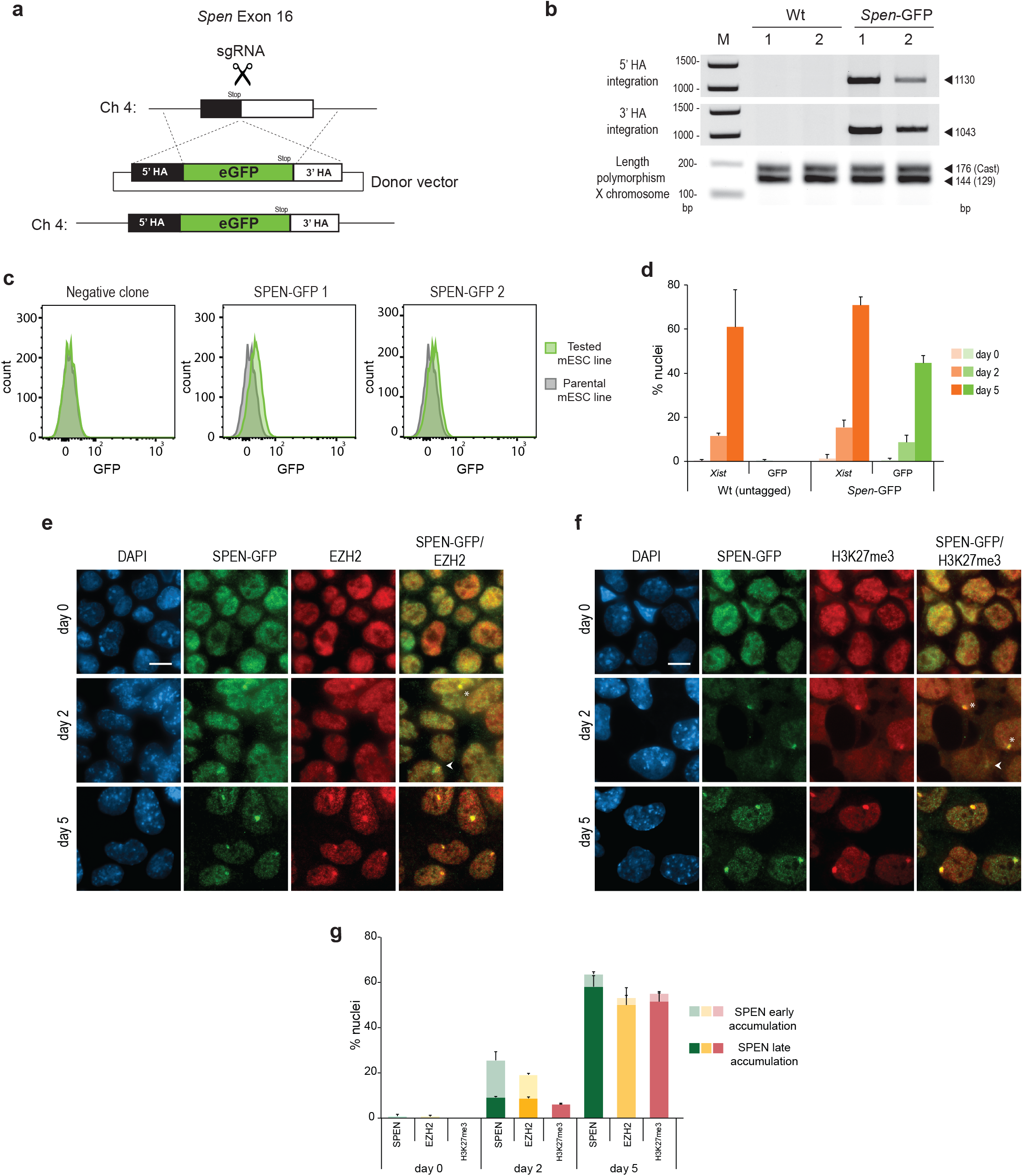
Generation and characterization of a SPEN-GFP C-terminal tag line. |Related to **Fig. 3**. **a** eGFP knock-in strategy in *Spen* exon 16 using the CRISPR/Cas9 system. **b** Genotyping of the *Spen*-GFP knock-in clones by PCR on gDNA to determine the specific integration of the 5’ and 3’ HA and the presence of two different X chromosomes making use of a length polymorphism. M = DNA ladder. **c** Flow cytometry histograms comparing the GFP fluorescence in the parental line (gray) with a negative clone **(left)** and two independent SPEN-GFP-tagged ESC clones **(middle and right)**. **d** Percentage of nuclei with *Xist* clouds and SPEN accumulation in Wt (untagged) and *Spen*-GFP ESCs. Quantification of **Fig. 3d.** Average percentage ± SD, n=2 independent clones, 200-500 nuclei quantified per replicate. **e** Double IF staining of SPEN-GFP (αGFP, green) and EZH2 (red) at day 0, 2 and 5 of differentiation of a *Spen*-GFP ESC line. Early accumulation is indicated with a white arrowhead, late accumulations with an asterisk. DNA is stained with DAPI (blue). Scale bar: 10 μm. **f** Double IF staining of SPEN-GFP (αGFP, green) and H3K27me3 (red) at day 0, 2 and 5 of differentiation of a *Spen*-GFP ESC line. Early accumulation is indicated with a white arrowhead, late accumulations with an asterisk. DNA is stained with DAPI (blue). Scale bar: 10 μm. **g** Quantification of **(e and f)**, showing the percentage of nuclei with early and late SPEN accumulations (green) and the percentage of nuclei with EZH2 (yellow) and H3K27me3 (light red) co-localizing with early or late SPEN accumulations. Average percentage ± SD, n=2 independent clones, 200-500 nuclei quantified per replicate.

**Supplementary Fig 5.**
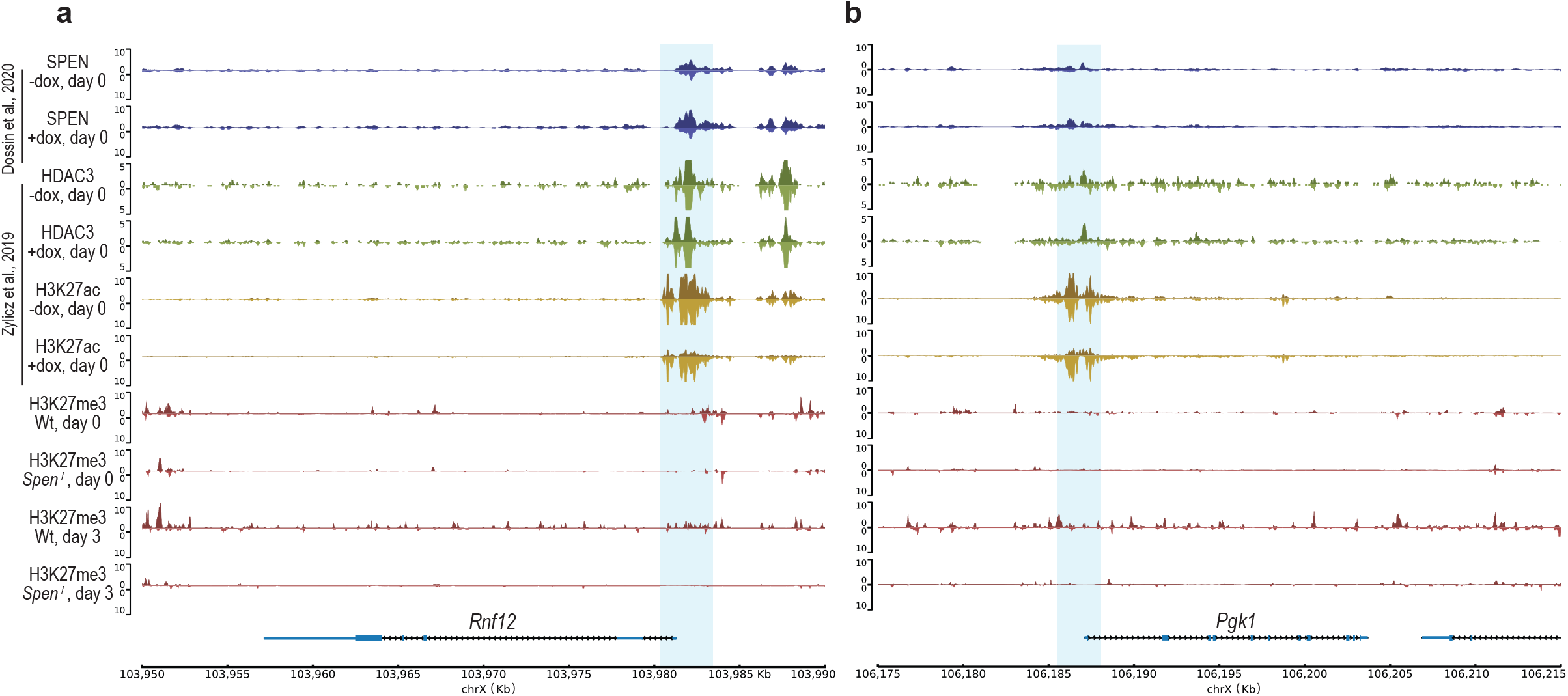
Chromatin features of X-linked genes. |Related to **Fig. 5**. **a-b** Genome browser tracks showing the allele-specific SPEN, HDAC3, H3K27ac and H3K27me3 binding for two X-linked genes: **a** *Rnf12* and **b** *Pgk1*. The top part (dark colour) of each track represents the Xi and the bottom (light colour) the Xa. SPEN CUT&RUN **(blue, top)** profile in *Xist* inducible undifferentiated ESCs (day 0) untreated or treated with doxycycline (24h)^25^. HDAC3 **(green, top-middle)** and H3K27ac ChIP-seq **(yellow, bottom-middle)** in *Xist* inducible undifferentiated ESCs (day 0) untreated or treated with doxycycline (24h)^30^. H3K27me3 ChIP-seq **(red, bottom)** in Wt and *Spen*^−/−^ ESCs at day 0 and 3 of monolayer differentiation. The light-blue square highlights the promoter region of *Rnf12* and *Pgk1.*

**Supplementary Fig 6.**
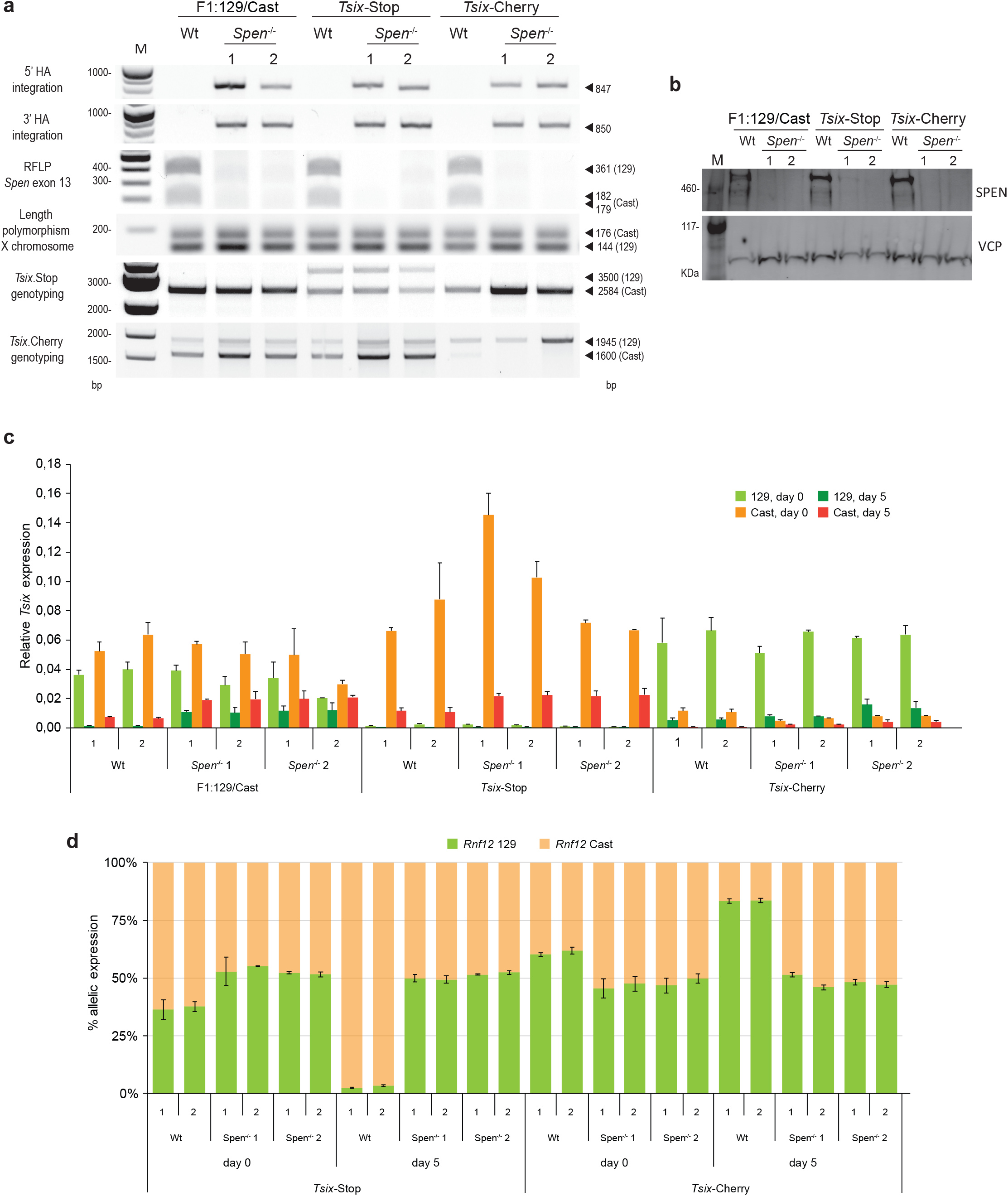
Generation and characterization of *Spen^−/−^* in *Tsix* defective ESC lines. | Related to **Fig. 5**. **a** Genotyping strategy to identify optimal *Spen*^−/−^ clones in F1:129/Cast, *Tsix-*Stop and *Tsix-*Cherry ESC lines, done by PCR on gDNA. Specific 5’ and 3’ integrations of the PuroR cassette in the *Spen* locus to identify *Spen* knockout lines. RFLP analysis on *Spen* exon 13 to determine the absence of the *Spen* 129 and/or Cast allele. Verification of the presence of two X chromosomes per ESC line, making use of a length polymorphism on the X chromosome. *Tsix*-Stop line genotyping, using a primer pair across the triple poly(A) signal blocking *Tsix* transcription. *Tsix*-Cherry line genotyping by determining the loss of the Cast band of a specific length polymorphism, indicating proper mCherry integration downstream of the *Tsix* promoter. M = DNA ladder. **b** SPEN western blot of Wt and *Spen*^−/−^ F1:129/Cast, *Tsix-*Stop and *Tsix-*Cherry ESC lines. VCP was used as a loading control. M = High molecular weight protein ladder. **c** Relative allele-specific *Tsix* expression at day 0 and 5 of monolayer differentiation of Wt and *Spen*^−/−^ F1:129/Cast, *Tsix-*Stop and *Tsix-*Cherry ESC lines, determined by RT-qPCR. Each individual clone was differentiated twice in biological duplicates. Average expression ± SD, n=2 biological replicates. **d** Percentage of *Rnf12* allelic expression at day 0 and 5 of monolayer differentiation of Wt and *Spen*^−/−^ F1:129/Cast, *Tsix-*Stop and *Tsix-*Cherry ESC lines, determined by RT-qPCR. Relative *Rnf12* allelic (129 and Cast) expression was normalized to *Rnf12* total expression and averaged ± SD, n=2 biological replicates.

## Methods

### Cell culture

Mouse ESCs were grown on male feeder cells and medium containing DMEM (Gibco), 15% Foetal Calf Serum (FCS), 0.1 mM non-essential amino acids (NEAA), 100 U mL^−1^ penicillin, 100 ug mL^−1^ streptomycin, 0.1 mM 2-mercaptoethanol (Gibco) and 1000 U mL^−1^ LIF. Previous to monolayer differentiation cells were plated in non-gelatinized plates to eliminate feeder cells. Then, plated at specific densities per time point in differentiation medium composed of IMDM-glutamax (Gibco), 15% FCS, 0.1 mM NEAA, 100 U mL^−1^ penicillin, 100 μg mL^−1^ streptomycin, 37.8 μL L^−1^ monothioglicerol and 50 mg mL^−1^ ascorbic acid. When appropriate, medium was supplemented with 2 μg mL^−1^ doxycycline.

### Gene editing using the CRISPR/Cas9 technology

Different female F1 2-1 hybrid (129/Sv-Cast/Ei) ESC lines with different genetic modifications to interrogate various aspects of XCI were targeted in this study (Supplementary Table 2). To generate *Spen* heterozygous and homozygous knockout clones two single-guide RNAs (sgRNA) targeting the 5’ (5’-AGTGCGCTTCGTCACTGCAC-3’) and 3’ (5’-TCCTCCCGCCCCGACGCGGA-3’) region of the *Spen* ORF were cloned in the Cas9-GFP pX458 vector (Addgene plasmid #48138). Compatible 5’ and 3’ Homology arms (HA) of approximately 500 bp were amplified by PCR from mouse gDNA and cloned in the pCR-Blunt-II-TOPO vector with a NdeI restriction site in-between the 5’ and 3’ HA. This site was used to insert a Puromycin resistance (PuroR) cassette flanked by loxP sites. The *ROSA26* locus was targeted using the pX458 vector coding for a sgRNA (5’-CGCCCATCTTCTAGAAAGAC-3’) compatible with the pFD46 expression vector ^25^, coding for the *Spen* cDNA and a hygromycin resistance cassette. To create an endogenous Spen C-terminal enhanced GFP (eGFP) knock-in, a sgRNA targeting the 3’ end of *Spen* ORF (5’-GATTGTCATTGCCTCGGTG-3’) was cloned in the Cas9-PuroR pX459 vector (Addgene plasmid #62988). The donor template was made using a gblock from Integrated DNA Technologies coding for compatible 5’ and 3’ HA of 600 bp with a NheI and AscI restrictions sites in-between the 5’ and 3’ HA, which were used to insert an eGFP in frame with the *Spen* coding sequence. The appropriate plasmid combinations were transfected into ESCs using lipofectamine 2000 and plated at low density to obtain single colonies, when appropriate medium was supplemented with 1 μg mL^−1^ puromycin (Sigma-Aldrich, P8833) for 48-72 hours or 250 μg mL^−1^ hygromycin B (Invitrogen, 10687010) for 7 days. Colonies were screened by PCR for correct integration of the desired construct (Supplementary Table 3). Positive clones were further characterized by western blot, RT-qPCR and/or FACS, and the presence of 2 X chromosomes and correct karyotype was also assessed.

### Protein extraction and western blot

To prepare nuclear extracts all procedures were done at 4°C and buffers supplemented with 1x protease inhibitors (Roche, 4693132001), 15 μM MG-132 (Sigma-Aldrich, C2211) and 0.5 mM DTT. Cells were harvested by scraping in cold PBS, collected and centrifuged (1500 rpm, 5 min, 4°C). The pellet was incubated in 5x times the pellet volume of Buffer A (10 mM Hepes pH 7.6, 1.5 mM MgCl2, 10 mM KCl) for 10 min, vortexed (30 sec) and centrifuged (3000 rpm, 5 min, 4°C). Then, the pellet was resuspended in 1,5x times Buffer C (20 mM Hepes pH 7.6, 25% glycerol, 420 mM NaCl, 1.5 mM MgCl2, 0.2 mM EDTA) and rotated (30 min, 4°C). The solution was centrifuged (14000 rpm, 10 min, 4°C) and the supernatant collected as nuclear extract. The protein concentration was measured using Nanodrop and all samples diluted to the same concentration using Buffer C. NuPAGE™ LDS Sample Buffer (4X) (Thermo Scientific, NP0007) containing 5% b-mercaptoethanol were added to nuclear extracts, and boiled (95°C, 5 min). For western blot analysis, the NuPAGE™ 3 to 8% Tris-Acetate gels (Invitrogen, EA03755) were used with Tris-Acetate SDS Running Buffer (pH 8.24) and the HiMark Pre-Stained Protein Standard (Invitrogen, LC5699). Wet transfer on a PVDF membrane was done overnight at 4 mA, with the NuPAGE Transfer Buffer (Invitrogen #NP00061), containing 10% methanol and 0.01% SDS. After blocking, the membrane was incubated with the appropriate antibodies: SPEN antibody (Abcam, ab290, 1:2000) and VCP antibody (Abcam, ab11433, 1:20000).

### RT-qPCR

Total RNA was isolated from cell pellets using the ReliaPrep RNA Cell Miniprep System (Promega, Z6012) and reversed transcribed using Superscript III (Invitrogen, 18080093) and random hexamers (Invitrogen, N8080127), following the manufacturer’s instructions. All RT-qPCRs were done using the GoTaq qPCR Master Mix (Promega, A6002) in a CFX384 real-time PCR detection system (Bio-Rad). Hist2h2aa1 was used as a normalization control, except in the RNA stability assay were β-actin was used. All expression primers are listed in Supplementary Table 4. The optimal allele-specific primer pair concentration to amplify the 129 and Cast allele at the same efficiency was optimized using pure 129, Cast and 129-Cast gDNA.

### RNA stability assay

Wt and *Spen*^−/−^ ESC lines with a doxycycline-responsive *Xist* promoter were cultured in medium supplemented with doxycycline to induce *Xist* expression for 4 days. While in doxycycline treatment, 5 μg mL^−1^ actinomycin D (Sigma-Aldrich, A1410) was added for different time (t= 0, 2, 4, 6 and 10 hours). Before collection, ESCs were plated in non-gelatinized plates to remove the feeders and harvested for RNA isolation, cDNA synthesis and RT-qPCR. Total *Xist* levels were determined and normalized to β-actin and the percentage of remaining *Xist* RNA was calculated by dividing *Xist* expression levels at the different times of collection relative to t=0. Using linear regression analysis the RNA decay rate constant (k decay) was calculated from the slope of the curve that best fitted our data and the RNA half-life (t1/2) obtained with the following formula t1/2 = ln2/(k decay).

### Immunofluorescence (IF)

Undifferentiated ESCs were attached to slides using a cytospin, while differentiating cells were grown on coverslips coated with gelatin or laminin. Cells were fixed with 4% paraformaldehyde (PFA) in PBS (10 min, RT), permeabilized with 0.4% Triton-X100 and 5% goat serum in PBS (15 min, on ice) and blocked with 10% goat serum in PBST, composed of 0.05% Tween-20 in PBS (30 min, RT). Primary antibodies incubation was done in blocking buffer (2h, RT), after three washings with PBST the slides were incubated with the secondary antibody (1h, RT). After three washings, the second washing containing DAPI, the slides were mounted with ProLong Gold antifade mounting medium (Invitrogen, P36930). The used primary antibodies were the following: αGFP (Abcam, ab290, 1:500), H3K27me2/3 (Active motif, 39535, 1:100) and EZH2(BD43 clone, kindly provided by Dr. Kristian Helin, 1:100). Images were taken using a ZEISS Axio Imager M2 including digital microscopy camera AxioCam 503 and analysed with ImageJ.

### IF combined with RNA Fluorescent in Situ Hybridization (FISH)

The *Xist* FISH probe was made from 2 μg of a 5.5 kb DNA fragment of mouse *Xist* comprising exons 3 to 7. Fluorescent labelling was done with dUTP SpectrumRed using the nick translation kit (Abbott, 07J00-001) overnight at 16°C. The probe was purified with ProbeQuant G-50 Micro Columns (GE, GE28-9034-08), combined with 100 μg mouse tRNA, 20 μg mouse Cot-1 DNA and 100 μg salmon sperm DNA and precipitated with 2 M NaAc (pH 5.6) and 100% EtOH. The pellet was resuspended in 50 μL hybridization mix (50% formamide, 10% dextran sulphate in 2xSSC) and stored at −20°C. Before use, 20 uL of probe was pre-hybridized (10 min, 75°C) with 0.5 μg mouse Cot-1 DNA supplemented with 10 mM vanadyl ribonucleoside complex (VRC) (NEB, S1402S) and 0.2 U μL^−1^ RNAseOUT (Invitrogen, 10777019).

Cells on coverslips or slides were fixed with 4% PFA in PBS (10 min, RT), permebilized with 0.5% Triton-X100 in PBS and blocked in TS-BSA buffer (0.1 M Tris-HCl (pH 7.5), 0.15 M NaCl, 2 mg mL^−1^ BSA (Jena Bioscience, BU-102) in H2O). The αGFP (Abcam, ab290, 1:500) primary antibody was incubated in blocking buffer (30 min, 37°C), washed in PBS (3x) and then incubated with the secondary antibody (30 min, 37°C). All solutions were supplemented with 10 μM VRC and 0.2 U μL^−1^ RNAseOUT. Slides were washed in PBS (3x), post-fixed with 4% PFA (10 min, RT) and washed again in PBS (x3). Dehydration was done with increasing ethanol concentration (70%, 90% and 100%). The pre-hybridized probe was added on top of the coverslips or slides (20h, 37°C, humid chamber). Various washings were done with 50% formamide/2x SSC (5 min, 37°C, 2x), 2x SSC (5 min, 37°C, 2x) and TS buffers (5 min, RT, 2x). Then the slides were mounted and visualized as explained in the IF section.

### FISH

The FISH only protocol was done as described in the IF-FISH section, skipping the IF section. Directly after the permeabilization, the dehydration step was performed.

### RNA-seq

Doxycycline treated cells were sorted to isolate dsRed positive cells upregulating *Xist* from the desired allele. RNA was isolated using the ReliaPrep RNA Cell Miniprep System. A total of 16 DNA libraries were created, according to the Smart-seq2 protocol ^49^, using the Nextera DNA Flex library prep kit (Illumina) to create a library from full-length cDNA. Samples were sequenced on a HiSeq2500 sequencer (50 bp single-end reads).

### Chromatin immunoprecipitation (ChIP)-seq

50×10^6 cells were trypsinised, resuspended and fixed in 50 mL warm medium and 1% PFA for 10 min at 37°C. 2.5 mL Glycine 2.5 M were added to the cells (final concentration 0.125 M) to quench the PFA, 5 min RT on a rotator. All buffers from now on contain protease inhibitors (Roche, 4693132001). Cells were washed twice with cold PBS. Then 1x in 10 mL Buffer 1 (10 mM Hepes pH 7.5, 10 mM EDTA, 0.5 mM EGTA, 0.75% Triton X-100) and 1x in Buffer 2 (10 mM Hepes pH 7.5, 200 mM NaCl, 1 mM EDTA, 0.5 mM EGTA), 10 min rotating at 4°C. Nuclei were then resuspended in Lysis/Sonication Buffer (150 mM NaCl, 25 mM Tris-HCl pH 7.5, 5 mM EDTA, 1% Triton, 0.1% SDS, 0.5% Sodium deoxycholate) and incubated 30 min on ice. Nuclei were then sonicated 2x 15 min (30’’ ON/OFF, max input, ice cold water) in a Bioruptor. A small fraction of the lysate was run on a gel to confirm size population of 100-500 bp. 1 μg of antibody (Cell Signaling, 9733S) was conjugated with 25 μL magnetic beads (Life Technologies, 10004D) for 3h rotating at 4°C. 25 ug of chromatin was IP’d with the Ab-beads overnight rotating at 4°C. Beads were washed 2x in standard RIPA buffer (140 mM NaCl, 10 mM Tris-HCl pH 7.5150 mM NaCl, 1 mM EDTA pH8.0, 0.5 mM EGTA pH 8.0, 1% Triton, 0.1% SDS, 0.5% Sodium deoxycholate), 1x in High Salt RIPA (same as standard RIPA but with 500 mM NaCl), 1x LiCl RIPA (same as standard RIPA but with 250 mM LiCl instead of NaCl) and rinsed once with TE, 10 min 4°C each wash. Chromatin was eluted with 450 μL of Elution Buffer (1% SDS; 0.1M NaHCO_3_ in H_2_O) with 22 μl protease K (10 mg mL^−1^) and 5ul RNAse A (10 mg mL^−1^) and shaken at 1000 rpm for 2 hours at 37°C first and then 65°C overnight. DNA was then Phenol-Chloroform extracted and resuspended in 20 μL H20. 0.25 μL was used per PCR to confirm the ChIP’s success. The concentration was then measured and a ChIP-seq library was prepared following the manufacturer’s instructions (ThruPLEX DNA, Takara Bio) and sequenced on an Illumina HiSeq 2500 sequencer (50 bp paired-end reads).

### NGS data analysis: allele-specific RNA-seq and ChIP-seq

Both the RNA-seq as the ChIP-seq data were processed allele-specifically. The single nucleotide polymorphism (SNPs) in the 129/Sv and Cast/Ei lines were downloaded from the Sanger institute (v.5 SNP142)^50^. These were used as input for SNPsplit v0.3.4^51^, to construct an N-masked reference genome based on mm10 in which all SNPs between 129/Sv and Cast/Ei were masked. The 50 bp single-end RNA-seq and 50 bp paired-end ChIP-seq reads were mapped to this N-masked reference genome using the default settings of hisat2 v2.2.1 and bowtie2 v2.4.1, respectively^52,53^. SNPsplit (--paired for the ChIP-seq analysis) was then used to assign the reads to either the 129/Sv or Cast/Ei bam file based on the best alignment or to a common bam file if mapping to a region without allele-specific SNPs. The allele-specific and unassigned bam files were sorted using samtools v1.10^54^.

For the RNA-seq, the number of mapped reads per gene were counted for both alleles separately using HTSeq v0.s12.4 (--nonunique=none -m intersection-nonempty)^55^ based on the gene annotation from ensembl v98. For each condition, genes with more than 20 allele-specific reads across both replicates were used to calculate the allelic ratio, defined as Xi/(Xi+Xa). For the day 0 and day 7, Cast/Ei and 129/Sv were used as the Xi and active allele (Xa), respectively, whereas for the day 3, Cast/Ei and 129/Sv were used as Xa and Xi, respectively. The allelic ratios of X-linked genes were visualized as violin plots with boxplots of the same data on top. Allelic ratios of individual genes were plotted along the X chromosome. Different conditions were plotted together showing only the X-linked genes that had at least 20 allele-specific reads in both conditions. We visualized differences in allelic ratios of X-linked genes between conditions by plotting both ratios on the different axes of a scatter plot. Genes were highlighted when they were identified as lowly silenced genes in *Spen*^−/−^ ESCs from^27^, which were defined as genes showing −0.05 < z < −0.2 where z (gene silencing) = [Xi/(Xi+Xa)]dox –[Xi/(Xi+Xa)]noDox. Significant differences between conditions were tested using a Mann-Whitney test with P-value < 0.05.

The allele-specific ChIP-seq bam files were normalized using the ‘callpeak’ and ‘bdgcmp’ functions of MACS2 v2.2.7.1^56^. We called broad peaks (-f BAMPE --broad --bdg) and used the Poisson P-value as method for normalizing the tracks. The input-normalized tracks were visualized using pyGenomeTracks v3.4.^57^. For validation, we downloaded several publicly available datasets. The SPEN CUT&RUN data (SRX5903674, SRX5903675, SRX5903676, SRX5903677, SRX5903678, SRX5903679, SRX5903682, SRX5903683)^25^, was processed similar to our analysis using a C57BL/6NJ-Cast/Ei reference genome. However, the allele-specific tracks were normalized based on the total number of mapped reads per sample. The scaling factor was calculated as 10^6 / total number of mapped reads and used as parameter --scaleFactor to both allelic tracks using deepTools bamCoverage v3.5.0.^58^. A binsize of 1 was used and paired-end reads were extended. The allele-specific tracks from HDAC3 and H3K27Ac (SRX4384412, SRX4384420, SRX4384476, SRX4384484, SRX4887836, SRX4887839) were downloaded from^30^. For all datasets, replicates for each condition were averaged using deepTools bigwigCompare v.3.5.0 with the settings ‘--operation mean --binSize 1’^58^. In the genome browser overview showing the allele-specific tracks, the y-axis was scaled for each group of samples separately.

## References

1. Lyon, M. F. Gene Action in the X-chromosome of the Mouse (Mus musculus L.). Nature 190, 372–373 (1961).

2. Borsani, G. et al. Characterization of a murine gene expressed from the inactive X chromosome. Nature 354, 56–58 (1991).

3. Brockdorff, N. et al. Conservation of position and exclusive expression of mouse Xist from the inactive X chromosome. Nature 351, 329–31 (1991).

4. Brockdorff, N. et al. The product of the mouse Xist gene is a 15 kb inactive X-specific transcript containing no conserved ORF and located in the nucleus. Cell 71, 515–526 (1992).

5. Lee, J. T., Davidow, L. S. & Warshawsky, D. Tsix, a gene antisense to Xist at the X-inactivation centre. Nat. Genet. 21, 400–404 (1999).

6. Stavropoulos, N., Lu, N. & Lee, J. T. A functional role for Tsix transcription in blocking Xist RNA accumulation but not in X-chromosome choice. Proc. Natl. Acad. Sci. U. S. A. 98, 10232–10237 (2001).

7. Luikenhuis, S., Wutz, A. & Jaenisch, R. Antisense Transcription through the Xist Locus Mediates Tsix Function in Embryonic Stem Cells. Mol. Cell. Biol. 21, 8512–8520 (2001).

8. Shibata, S. & Lee, J. T. Tsix Transcription- versus RNA-Based Mechanisms in Xist Repression and Epigenetic Choice. Curr. Biol. 14, 1747–1754 (2004).

9. Sado, T., Hoki, Y. & Sasaki, H. Tsix silences Xist through modification of chromatin structure. Dev. Cell 9, 159–165 (2005).

10. Navarro, P., Pichard, S., Ciaudo, C., Avner, P. & Rougeulle, C. Tsix transcription across the Xist gene alters chromatin conformation without affecting Xist transcription: Implications for X-chromosome inactivation. Genes Dev. 19, 1474–1484 (2005).

11. Sun, B. K., Deaton, A. M. & Lee, J. T. A transient heterochromatic state in Xist preempts X inactivation choice without RNA stabilization. Mol. Cell 21, 617–628 (2006).

12. Mutzel, V. et al. A symmetric toggle switch explains the onset of random X inactivation in different mammals. Nat. Struct. Mol. Biol. (2019). doi:10.1038/s41594-019-0214-1

13. Mutzel, V. & Schulz, E. G. Dosage Sensing, Threshold Responses, and Epigenetic Memory: A Systems Biology Perspective on Random X-Chromosome Inactivation. BioEssays 42, 1–14 (2020).

14. Loda, A. & Heard, E. Xist RNA in action: Past, present, and future. PLoS Genet. 15, 1–17 (2019).

15. Brockdorff, N. & Turner, B. M. Dosage compensation in mammals. Cold Spring Harb. Perspect. Biol. 7, (2015).

16. McHugh, C. A. et al. The Xist lncRNA interacts directly with SHARP to silence transcription through HDAC3. Nature 521, 232–236 (2015).

17. Chu, C. et al. Systematic discovery of Xist RNA binding proteins. Cell 161, 404–416 (2015).

18. Minajigi, A. et al. A comprehensive Xist interactome reveals cohesin repulsion and an RNA-directed chromosome conformation. Science (80-.). 316, (2015).

19. Monfort, A. et al. Identification of Spen as a crucial factor for Xist function through forward genetic screening in haploid embryonic stem cells. Cell Rep. 12, 554–561 (2015).

20. Moindrot, B. et al. A Pooled shRNA Screen Identifies Rbm15, Spen, and Wtap as Factors Required for Xist RNA-Mediated Silencing. Cell Rep. 12, 562–572 (2015).

21. Ariyoshi, M. & Schwabe, J. W. R. A conserved structural motif reveals the essential transcriptional repression function of spen proteins and their role in developmental signaling. Genes Dev. 17, 1909–1920 (2003).

22. Oswald, F. et al. A phospho-dependent mechanism involving NCoR and KMT2D controls a permissive chromatin state at Notch target genes. Nucleic Acids Res. 44, 4703–4720 (2016).

23. Shi, Y. et al. Sharp, an inducible cofactor that integrates nuclear receptor repression and activation. Genes Dev. 15, 1140–1151 (2001).

24. Oswald, F. et al. SHARP is a novel component of the Notch/RBP-Jκ signalling pathway. EMBO J. 21, 5417–5426 (2002).

25. Dossin, F. et al. SPEN integrates transcriptional and epigenetic control of X-inactivation. Nature 578, 455–460 (2020).

26. Loda, A. et al. Genetic and epigenetic features direct differential efficiency of Xist-mediated silencing at X-chromosomal and autosomal locations. Nat. Commun. 8, (2017).

27. Nesterova, T. B. et al. Systematic allelic analysis defines the interplay of key pathways in X chromosome inactivation. Nat. Commun. 10, 1–15 (2019).

28. Wutz, A. & Jaenisch, R. A shift from reversible to irreversible X inactivation is triggered during ES cell differentiation. Mol. Cell 5, 695–705 (2000).

29. Ogawa, Y. & Lee, J. T. Xite, X-Inactivation Intergenic Transcription Elements that Regulate the Probability of Choice. 11, 731–743 (2003).

30. Żylicz, J. J. et al. The Implication of Early Chromatin Changes in X Chromosome Inactivation. Cell 176, 182–197.e23 (2019).

31. Jonkers, I. et al. RNF12 Is an X-Encoded Dose-Dependent Activator of X Chromosome Inactivation. Cell 139, 999–1011 (2009).

32. Heard, E. et al. Methylation of histone H3 at Lys-9 Is an early mark on the X chromosome during X inactivation. Cell 107, 727–738 (2001).

33. Rougeulle, C. et al. Differential Histone H3 Lys-9 and Lys-27 Methylation Profiles on the X Chromosome. Mol. Cell. Biol. 24, 5475–5484 (2004).

34. Loos, F. et al. Xist and Tsix Transcription Dynamics Is Regulated by the X-to-Autosome Ratio and Semistable Transcriptional States. Mol. Cell. Biol. 36, 2656–2667 (2016).

35. Barakat, T. S., Rentmeester, E., Sleutels, F., Grootegoed, J. A. & Gribnau, J. Precise BAC targeting of genetically polymorphic mouse ES cells. Nucleic Acids Res. 39, 6–13 (2011).

36. Carter, A. C. et al. Spen links rna-mediated endogenous retrovirus silencing and x chromosome inactivation. Elife 9, 1–58 (2020).

37. Kuroda, K. et al. Regulation of marginal zone B cell development by MINT, a suppressor of Notch/RBP-J signaling pathway. Immunity 18, 301–312 (2003).

38. Hoki, Y. et al. A proximal conserved repeat in the Xist gene is essential as a genomic element for X-inactivation in mouse. Development 136, 139–146 (2009).

39. Engreitz, J. M. et al. The Xist lncRNA exploits three-dimensional genome architecture to spread across the X chromosome. Science (80-.). 341, 1237973 (2013).

40. Lee, J. T. & Lu, N. Targeted mutagenesis of Tsix leads to nonrandom X inactivation. Cell 99, 47–57 (1999).

41. Xu, N., Tsai, C. L. & Lee, J. T. Transient homologous chromosome pairing marks the onset of X inactivation. Science (80-.). 311, 1149–1152 (2006).

42. Aeby, E. et al. Decapping enzyme 1A breaks X-chromosome symmetry by controlling Tsix elongation and RNA turnover. Nat. Cell Biol. 22, 1116–1129 (2020).

43. Monkhorst, K., Jonkers, I., Rentmeester, E., Grosveld, F. & Gribnau, J. X Inactivation Counting and Choice Is a Stochastic Process: Evidence for Involvement of an X-Linked Activator. Cell 132, 410–421 (2008).

44. Gayen, S. et al. Article A Primary Role for the Tsix lncRNA in Maintaining Article A Primary Role for the Tsix lncRNA in Maintaining Random X-Chromosome Inactivation. CellReports 11, 1251–1265 (2015).

45. Galupa, R. et al. A Conserved Noncoding Locus Regulates Random Monoallelic Xist Expression across a Topological Boundary. Mol. Cell 77, 352–367.e8 (2020).

46. Donohoe, M. E., Silva, S. S., Pinter, S. F., Xu, N. & Lee, J. T. The pluripotency factor Oct4 interacts with Ctcf and also controls X-chromosome pairing and counting. Nature 460, 128–132 (2009).

47. Navarro, P. et al. Molecular coupling of Tsix regulation and pluripotency. Nature 468, 457–460 (2010).

48. Gontan, C. et al. RNF12 initiates X-chromosome inactivation by targeting REX1 for degradation. Nature 485, 386–390 (2012).

49. Picelli, S. et al. Smart-seq2 for sensitive full-length transcriptome profiling in single cells. Nat. Methods 10, 1096–1100 (2013).

50. Keane, T. M. et al. Mouse genomic variation and its effect on phenotypes and gene regulation. Nature 477, 289–294 (2011).

51. Krueger, F. & Andrews, S. R. SNPsplit: Allele-specific splitting of alignments between genomes with known SNP genotypes. F1000Research 5, 1479 (2016).

52. Kim, D., Langmead, B. & Salzberg, S. L. HISAT: A fast spliced aligner with low memory requirements. Nat. Methods 12, 357–360 (2015).

53. Langmead, B. & Salzberg, S. L. Fast gapped-read alignment with Bowtie 2. Nat. Methods 9, 357–359 (2012).

54. Li, H. et al. The Sequence Alignment/Map format and SAMtools. Bioinformatics 25, 2078–2079 (2009).

55. Anders, S., Pyl, P. T. & Huber, W. HTSeq-A Python framework to work with high-throughput sequencing data. Bioinformatics 31, 166–169 (2015).

56. Zhang, Y. et al. Model-based analysis of ChIP-Seq (MACS). Genome Biol. 9, (2008).

57. Lopez-Delisle, L. et al. pyGenomeTracks: reproducible plots for multivariate genomic data sets. Bioinformatics 1–2 (2020). doi:10.1093/bioinformatics/btaa692

58. Ramírez, F. et al. deepTools2: a next generation web server for deep-sequencing data analysis. Nucleic Acids Res. 44, W160–W165 (2016).

