## Supplementary Tables 1 to 4 for "SPEN is Required for *Xist* Upregulation during Initiation of X Chromosome Inactivation"

**Supplementary Table 1.** Overview of publications that studied *Spen* in relation to XCI. KO = knockout, KD = knockdown, n.d. = not-determined.

| **Reference** | **Doxycycline inducible *Xist* or physiological XCI** | ***Spen* KO or KD** | ***Xist* upregulation** | **XCI** |
|---|---|---|---|---|
| [1] | Doxycycline inducible *Xist*/ physiological XCI | KD, siRNA | + | - |
| [2] | Doxycycline inducible *Xist* | KD, siRNA | + | - |
| [3] | n.d. | n.d. | n.d. | n.d. |
| [4] | Doxycycline inducible *Xist*/ physiological XCI | KD, shRNA | + | - |
| [5] | Doxycycline inducible *Xist* | KO | + | - |
| [6] | Doxycycline inducible *Xist* | KO | + | - |
| [7] | Doxycycline inducible *Xist* | KD, Auxin-inducible degron | + | - |
| [8] | Doxycycline inducible *Xist*/ physiological XCI | KO | + / ? | - |
| [9] | Doxycycline inducible *Xist* | KO | + | - |
| This study | Doxycycline inducible *Xist*/ physiological XCI | KO | +/- | - |

**Supplementary Table 2.** Genetically modified ESC lines generated in this study.

| Genetic modification | Donor vector, sgRNA(s) | Transfected ESC line | ESC line source | Genotype | Clone names | Used in |
|---|---|---|---|---|---|---|
| *Spen* homozygous and heterozygous knockout | 5'HA – PuroR – 3'HA, 5'+ 3' sgRNA | Doxycycline responsive endogenous *Xist* promoter (Clone 87) | [10] | Wt | Wt 1: parental<br>Wt2: C1 (transfected, no deletion) | Figure 1 and 4 |
| | | | | *Spen*$^{+/-}$ | *Spen*$^{+/-}$ 1: D7<br>*Spen*$^{+/-}$ 2: E7 | |
| | | | | *Spen*$^{-/-}$ | *Spen*$^{-/-}$ 1: B3<br>*Spen*$^{-/-}$ 2: G4 | |
| | | F1:129/Cast | [11] | Wt | Parental | Figure 5 |
| | | | | *Spen*$^{-/-}$ | *Spen*$^{-/-}$ 1: A8<br>*Spen*$^{-/-}$ 2: A10 | |
| | | *Tsix*-Stop | [12] | Wt | Parental | |
| | | | | *Spen*$^{-/-}$ | *Spen*$^{-/-}$ 1: A5<br>*Spen*$^{-/-}$ 2: B6 | |
| | | *Tsix*-Cherry | [13] | Wt | Parental | |
| | | | | *Spen*$^{-/-}$ | *Spen*$^{-/-}$ 1: A4<br>*Spen*$^{-/-}$ 2: H2 | |
| *Spen* cDNA rescue in *Spen* homozygous knockout line | pFD46 vector [7] | *Spen*$^{-/-}$ (Clone B3) | This study | Wt | Clone 87 | Figure 2 |
| | | | | *Spen*$^{-/-}$ | parental | |
| | | | | *Spen*$^{-/-}$ + *Spen* cDNA in *ROSA26* locus | Clone A: B6<br>Clone B: E4<br>Clone C: F3 | |
| *Spen* C-terminal eGFP tag | 5'HA – eGFP – 3'HA, 3' sgRNA | Doxycycline responsive endogenous *Xist* promoter (Clone 87) | [10] | Wt | Wt1: parental<br>Wt2: G5 (transfected, no integration) | Figure 3 |
| | | | | *Spen*-eGFP | Spen-GFP 1: B2<br>Spen-GFP 2: D7 | |

**Supplementary Table 3.** List of genotyping primers.

| **Name** | **Sequence** | **Source** |
|---|---|---|
| 9.5’-PuroR.integration.Fw | GTGTCCCTCATGCAAAGTGG | |
| 12.5’-PuroR.integration.Rv | TTAATTGTAGCCGCGTTCTAAC | |
| 14.3’-PuroR.integration.Fw | AGACTGCCTTGGGAAAAGCG | |
| 16.3’-PuroR.integration.Rv | CTGTACCCGAAGCACCATTT | |
| 250.RFLP1.Exon13.BlpI.Fw | GACTTCAGCGTGAGGCAGAG | |
| 250.RFLP1.Exon13.BlpI.Rv | CACGCAGCCTATACCACCTG | |
| X-LP–Dxmit65.Fw | ATATTAAGGGAGGTAACAAAGACCC | Whitehead Institute at MIT; Center for Genome Research |
| X-LP–Dxmit65.Rv | GGTTTCTGTGATTGCTATAGGACA | |
| 2.Across.Xist.Dox.Promoter.Fw | CCCAGATGGGCAAGTTTAGA | |
| 4.Across.Xist.Dox.Promoter.Rv | CAGGACATCTGGGGCTATACA | |
| 48.Spen.Rosa26.integration.Right.Fw | TTTGCATTCCAAAAGGAACC | |
| 49.Spen.Rosa26.integration.Right.Rv | ATACGAGGTCGCCAACATCT | |
| 55.SpenRRM1.Fw | CGCTCCCTGTTATCTGAAGC | |
| 56.Spen.Flag.Rv | AAGGACCACGACGGAGACTA | |
| 1.3’-eGFP.integration.Fw | TCCTTGAAGTCGATGCCCTT | |
| 2.3’-eGFP.integration.Rv | GTGGAAACCGACTACTGCCT | |
| 4.5’-eGFP.integration.Fw | GTTCTGCACATCCGACCAAG | |
| 7.5’-eGFP.integration.Rv | CACATGAAGCAGCACGACTT | |
| 350-36. Fw Tsix LP, for DNA | AGTGCAGCGCTTGTGTCA | [13] |
| 351-41. Rv Tsix LP, for DNA | TATTACCCACGCCAGGCTTA | |
| 356.Tsix Stop genotyping - 3F | CTTTGGTTTTGATGCGGATT | |
| 357.Tsix Stop genotyping - 3R | GCCTCTGTCACTCCATCTCC | |

**Supplementary Table 4.** List of expression and allele-specific primers.

| Name | Sequence | Source |
|---|---|---|
| Xist_129_F4 | GGAAGAAGGTAGGATTCTACCTCTTC | |
| Xist_cast_F4 | GGAAGAAGGTAGGATTCTACCTCATG | |
| Xist_R4 | GCCAGCACTGATCTCAAGC | |
| 358.Xist_Ex1-2_F | GGATCCTGCTTGAACTACTGC | [14] |
| 359.Xist_Ex1-2_R | CAGGCAATCCTTCTTCTTGAG | |
| rnf12_Ex5_129F | CAGAACGGGAAAAGGTACGC | |
| rnf12_Ex5_129R | ATACCGGCAGAGAGATAGTATAGCTTGC | |
| rnf12_Ex5_Cast_F | TTCAGAACGGGAAAAGGTACGT | |
| rnf12_Ex5_Cast_R | GAATACCGGCAGAGAGATAGTATAGCTTGT | |
| Tsix_In3_129_R | CAGGGCTACCCTGGAAAACT | |
| Tsix_In3_Cast_R | CCAGGGCTATCCTGGAAAATC | |
| Tsix_In3_F | TGCATTAGCTGCTCCTCCTT | |
| 297.Hist2h3c1.qPCR.Ctl.Fw | GTTTGCGCTTTCGTGATGTC | |
| 298.Hist2h3c1.qPCR.Ctl.Rv | CCCCACCGGGAACTGTAG | |
| 295.Rex1_F | CTAAAGCAAGACGAGGCAAG | [15] |
| 296.Rex1_R | AGAATGGGTTCGGAAAACTC | |
| Nanog qPCR_FW | AGGATGAAGTGCAAGCGGTG | [16] |
| Nanog qPCR_RV | TGCTGAGCCCTTCTGAATCAG | |
| Gata6.Fw | GAGCTGGTGCTACCAAGAGG | [17] |
| Gata6.Rv | TGCAAAAGCCCATCTCTTCT | |
| 372_ß-Actin qPCR_FW | ACTATTGGCAACGAGCGGTTC | [18] |
| 373_ ß-Actin qPCR_RV | AGAGGTCTTTACGGATGTCAACG | |
| 94. SPEN Exon10. qPCR Fw | GCAAATCGGGAAAGCCAACT | [5] |
| 95. SPEN Exon11. qPCR Rv | CTGCACTCCAGTCTTCATGC | |

### References (Supplementary Tables)


1. Chu, C. *et al.* Systematic discovery of Xist RNA binding proteins. *Cell* **161**, 404–416 (2015).

2. McHugh, C. A. *et al.* The Xist lncRNA interacts directly with SHARP to silence transcription through HDAC3. *Nature* **521**, 232–236 (2015).

3. Minajigi, A. *et al.* A comprehensive Xist interactome reveals cohesin repulsion and an RNA-directed chromosome conformation. *Science (80-. ).* **316**, (2015).

4. Moindrot, B. *et al.* A Pooled shRNA Screen Identifies Rbm15, Spen, and Wtap as Factors Required for Xist RNA-Mediated Silencing. *Cell Rep.* **12**, 562–572 (2015).

5. Monfort, A. *et al.* Identification of Spen as a crucial factor for Xist function through forward genetic screening in haploid embryonic stem cells. *Cell Rep.* **12**, 554–561 (2015).

6. Nesterova, T. B. *et al.* Systematic allelic analysis defines the interplay of key pathways in X chromosome inactivation. *Nat. Commun.* **10**, 1–15 (2019).

7. Dossin, F. *et al.* SPEN integrates transcriptional and epigenetic control of X-inactivation. *Nature* **578**, 455–460 (2020).

8. Carter, A. C. *et al.* Spen links rna-mediated endogenous retrovirus silencing and x chromosome inactivation. *Elife* **9**, 1–58 (2020).

9. Trotman, J. B. *et al.* Elements at the 5' end of Xist harbor SPEN-independent transcriptional antiterminator activity. *Nucleic Acids Res.* 1–18 (2020). doi:10.1093/nar/gkaa789

10. Loda, A. *et al.* Genetic and epigenetic features direct differential efficiency of Xist-mediated silencing at X-chromosomal and autosomal locations. *Nat. Commun.* **8**, (2017).

11. Barakat, T. S., Rentmeester, E., Sleutels, F., Grootegoed, J. A. & Gribnau, J. Precise BAC targeting of genetically polymorphic mouse ES cells. *Nucleic Acids Res.* **39**, 6–13 (2011).

12. Luikenhuis, S., Wutz, A. & Jaenisch, R. Antisense Transcription through the Xist Locus Mediates Tsix Function in Embryonic Stem Cells. *Mol. Cell. Biol.* **21**, 8512–8520 (2001).

13. Loos, F. *et al.* Xist and Tsix Transcription Dynamics Is Regulated by the X-to-Autosome Ratio and Semistable Transcriptional States. *Mol. Cell. Biol.* **36**, 2656–2667 (2016).

14. Chureau, C. *et al.* Ftx is a non-coding RNA which affects Xist expression and chromatin structure within the X-inactivation center region. *Hum. Mol. Genet.* **20**, 705–718 (2011).

15. Gontan, C. *et al.* RNF12 initiates X-chromosome inactivation by targeting REX1 for degradation. *Nature* **485**, 386–390 (2012).

16. Navarro, P. *et al.* Molecular Coupling of Xist Regulation and Pluripotency. **321**, 1693–1696 (2008).

17. Shimosato, D., Shiki, M. & Niwa, H. Extra-embryonic endoderm cells derived from ES cells induced by GATA Factors acquire the character of XEN cells. **12**, 1–12 (2007).

18. Jonkers, I. *et al.* Xist RNA Is Confined to the Nuclear Territory of the Silenced X Chromosome throughout the Cell Cycle. *Mol. Cell. Biol.* **28**, 5583–5594 (2008).
